# Larval diet affects adult reproduction but not survival regardless of injury and infection stress in *Drosophila melanogaster*

**DOI:** 10.1101/2020.10.16.342618

**Authors:** Eevi Savola, Pedro Vale, Craig Walling

## Abstract

Early-life conditions have profound effects on many life-history traits. In particular, early-life diet affects both juvenile development, and adult survival and reproduction. Early-life diet also has consequences for the ability of adults to withstand stressors such as starvation, temperature and desiccation. However, it is less well known how early-life diet influences the ability of adults to respond to infection. Here we test whether varying the larval diet of female *Drosophila melanogaster* (through altering protein to carbohydrate ratio, P:C) influences the long-term response to injury and infection with the bacterial pathogen *Pseudomonas entomophila*. Given previous work manipulating adult dietary P:C, we predicted that adults from larvae raised on higher P:C diets would be more likely to survive infection and have increased reproduction, but shorter lifespans and an increased rate of ageing. For larval development, we predicted that low P:C would lead to a longer development time and lower viability. We found that early-life and lifetime egg production were highest at intermediate to high larval P:C diets, but there was no effect of larval P:C on adult survival. Larval diet had no effect on survival or reproduction post-infection. Larval development was quickest on intermediate P:C and egg-to-pupae and egg-to-adult viability were higher on higher P:C. Overall, despite larval P:C affecting several traits measured in this study, we saw no evidence that larval P:C altered the consequence of infection or injury for adult survival and early-life and lifetime reproduction. Taken together, these data suggest that larval diets appear to have a limited impact on adult response to infection.

## INTRODUCTION

Early-life conditions are important in determining many key life-history traits (reviewed in Metcalfe and Monaghan, 2001, 2003). In particular, diet in early-life has been shown to have profound effects on later life-history traits such as survival and reproduction, and poor early nutrition can have costs associated with catch-up growth in adulthood (reviewed in Metcalfe and Monaghan, 2001, 2003). Nutrition is also important for the ability of an organism to respond to a number of key environmental stresses such as infection or temperature stress, as has been demonstrated in both juveniles (e.g. Lee *et al.* 2006; Venesky *et al.* 2012; Kutz, Sgrò and Mirth, 2019) and adults (e.g. Peck, Babcock and Alexander, 1992; Kim, Jang and Lee, 2020; Ponton *et al.* 2020). However, work on the effect of early-life diet on adult responses to environmental stress is much more limited (but see e.g. Andersen et al. 2010; Kelly and Tawes 2013; Knutie et al. 2017). To investigate the long-term effects of early-life diet on adult traits and infection stress resistance, here we combine multiple larval diets, apply injury and infection to the adults and measure both larval and adult life-history responses in *Drosophila melanogaster.*

A vast literature exists using various approaches to manipulate diet and investigate the consequences of these manipulations (reviewed in Simpson and Raubenheimer, 2012). A particularly well-investigated manipulation is adult dietary restriction (DR), the restriction of calories or a particular nutrient without malnutrition, which has been shown to increase lifespan, delay ageing and reduce reproduction across a wide range of species (e.g. Mair and Dillin, 2008; Simpson *et al.* 2017). Recent evidence suggests that this effect is mostly driven by changes in the protein to non-protein ratio of the diet, often protein to carbohydrate (P:C) ratios, particularly in insects (e.g. Lee *et al.* 2008; Simpson *et al.* 2017, but see Speakman, Mitchell and Mazidi, 2016). Regarding the effects of juvenile diet on juvenile and adult traits, there have been many studies testing the effects of caloric content (e.g. May, Doroszuk and Zwaan, 2015; Adler, Telford and Bonduriansky, 2016; House *et al.* 2016; Littlefair and Knell, 2016; Hooper *et al.* 2017; Krittika, Lenka and Yadav, 2019). As it has become clearer that macronutrient content is more important than total caloric content, recent work has shifted to testing how the macronutrient composition of the juvenile diet may affect both juvenile and adult traits (reviewed in Nestel *et al.* 2016). However, these studies often do not consider additional stressors (but see e.g. Andersen *et al.* 2010; Kelly and Tawes, 2013; Pascacio-Villafán *et al.* 2016).

Changing juvenile diet has been shown to alter the rate and success of the developmental period in both holometabolous (reviewed in Nestel *et al.* 2016) and hemimetabolous insects (e.g. Hunt *et al.* 2004; Kelly and Tawes, 2013; Houslay *et al.* 2015). In general, juveniles on higher or intermediate P:C diets have a quicker development rate and improved development success (e.g. Matavelli *et al.* 2015; Rodrigues *et al.* 2015; Silva-Soares *et al.* 2017, but see Cordes *et al.* 2015; Houslay *et al.* 2015; Davies *et al.* 2018; Gray, Simpson and Polak, 2018; Kim *et al.* 2019). In holometabolous insects, larvae have to pass several size assessment thresholds for successful pupation, and it has been suggested larvae feed until they have enough resources for metamorphosis and to survive the non-feeding state of pupation (reviewed in Mirth and Riddiford, 2007; Nestel *et al.* 2016). As amino acids from protein in the diet signal a cell cycle for growth of tissues (Britton and Edgar 1998; Colombani et al. 2003), and larvae do not develop on diets lacking in essential amino acids (e.g. Chang, 2004), it seems that higher larval P:C diets facilitate quicker growth and accumulation of essential resources that allow successful development into adulthood. There may be an upper limit after which increasing P:C has detrimental effects, potentially due to toxic effects of protein metabolism (Fanson et al. 2012), for example the accumulation of toxic wastes in food (reviewed in Simpson and Raubenheimer, 2009) or the highest P:C diets being limiting in carbohydrates, but the exact reasons are currently unknown.

Early-life diet has also been shown to have important consequences for many adult life-history traits, including reproduction, lifespan and ageing (reviewed in Metcalfe and Monaghan, 2001). In insects, measures of both early-life and lifetime egg production peak on higher or intermediate larval P:C diets (e.g. Rodrigues *et al.* 2015; Silva-Soares *et al.* 2017; Duxbury and Chapman, 2019, but see Matavelli *et al.* 2015). For lifespan, results of early-life dietary manipulation in insects are mixed, with lifespan being maximised at different P:C levels, and even no effect of P:C depending on the study (e.g. Runagall-McNaull, Bonduriansky and Crean, 2015; Stefana *et al.* 2017; Davies *et al.* 2018; Duxbury and Chapman, 2019). Indeed, a recent meta-analysis showed no consistent effect of early-life diet on adult lifespan across taxa (English and Uller 2016). The age-related decline in various traits may also be altered by larval diet, however the direction of the effect is again unclear, with higher P:C or calorie diets leading to quicker, slower or having no effect on ageing (Tu and Tatar 2003; May et al. 2015; Adler et al. 2016; Hooper et al. 2017). The effect of larval diet on adult reproduction may be a result of adults being able to use nutrient stores of, for example, protein or lipids in body tissues, including the fat body and haemolymph (reviewed in Boggs, 2009; Nestel *et al.* 2016). However, it is less clear how these stored resources could affect lifespan. Potential explanations for inconsistencies in results across studies and life-history traits include that stored nutrients may trade-off between different adult life-history traits in an environment or species-specific manner, that juvenile diet effects may be dependent on adult food environment, or that storage of nutrients can be re-allocated in adulthood, for example by reabsorption of flight muscles (reviewed in Boggs, 2009; Nestel *et al.* 2016). Overall, it seems that increasing P:C in the larval diet increases reproduction and juvenile diet often has effects on adult lifespan, but the directionality of the effects are inconsistent.

Despite the wealth of information on how larval diet affects multiple adult traits, studies focusing on adult stress resistance are rarer, despite the likelihood that stress resistance is a key trait in natural populations (e.g. Hoffman and Hercus, 2000; Van Voorhies, Fuchs and Thomas, 2005; Kawasaki *et al.* 2008; Adamo, 2020). Some data exist on a small number of environmental stressors including temperature, desiccation and starvation (Andersen et al. 2010; Pascacio-Villafán et al. 2016; Davies et al. 2018), however the direction of effects are often mixed and potentially stress-specific. Particularly poorly studied is the effect of larval diet on adult infection response. In *Anopheles gambiae,* melanising ability decreased with severity of larval calorie restriction (Suwanchaichinda and Paskewitz 1998). Female *Gryllus texensis* crickets on lower P:C as nymphs and adults survive better over five days post-infection (Kelly and Tawes 2013). As this study changed both adult and larval diet, it is not possible to disentangle the effect of larval diet alone. Without a direct immune stress, there is evidence for differential adult response to immune challenge due to larval diet. For example, the production of antimicrobial peptides (AMPs) decreased with lower larval P:C in *D. melanogaster* (Fellous and Lazzaro 2010). For other immune response measures, in *Lestes viridis* damselflies, lower calories and starvation led to reduced phenoloxidase (PO) activity and haemocyte numbers/levels in adults (Rolff et al. 2004; De Block and Stoks 2008). However, to our knowledge, no study to date has tested the effect of larval dietary P:C on adult life-history traits when exposed to infection or injury stress.

Several hypotheses have been put forward to explain the effect of larval diet on adult survival post-infection. These include increased stress response capability due to overall better body condition, and increased investment into immunity, either through the growth of specific tissues, or through increased availability of limiting nutrients (Fellous and Lazzaro 2010). The first suggestion is supported by studies where both immune response and body condition are lower with starvation (Suwanchaichinda and Paskewitz 1998; Rolff et al. 2004), but the independent effects are difficult to separate (Fellous and Lazzaro 2010). The second hypothesis is supported by studies where indicators of immune response increase in adults or pupae outside of effects on general body condition with a diet higher in P:C in *D. melanogaster* (Fellous and Lazzaro 2010) or plant diets of worse quality in *Epirrita autumnata* moths (Klemola et al. 2007). In general protein seems to be an important nutrient in relation to survival post-infection (e.g. Lee *et al.* 2006; Povey *et al.* 2009, 2014; Cotter *et al.* 2011, 2019; Savola *et al.* 2020), suggesting individuals developing on higher larval P:C diets should have improved resistance to infection.

To test the effects of larval P:C on larval and adult life-history and adult survival post-infection, we reared larvae on various P:C diets and exposed female adults to injury and infection stress (with a bacterial pathogen, *Pseudomonas entomophila*). For the larvae, we measured development time to adulthood and measures of viability (egg-to-pupae, egg-to-adult and pupae-to-adult viability). For the adults, we measured the key life-history traits of survival and reproduction. We predicted that low P:C larval food would lead to longer development time and lower viability across all stages. If larval diet affects adult life-history traits independent of adult food, we predicted a similar effect to that observed when P:C ratio is manipulated in adults (e.g. Lee *et al.* 2008; Jensen *et al.* 2015; Savola *et al.* 2020), with low larval P:C extending lifespan, and reducing reproduction and the senescent decline in egg laying. Conversely, if larval diet has no long-term effects on life-history traits, we would expect to see similar survival and reproduction patterns across all diets. As low P:C diets have been found to be especially detrimental for survival post-infection in our host-pathogen system (Savola et al. 2020), we predicted that low larval P:C would reduce survival and reproduction to a greater extent in injured and infected flies than in control flies.

## METHODS

### LARVAL DIETS

Larval diets consisted of five diets varying in P:C composition from 1:16 to 2:1 P:C (corresponding to 5 to 61% protein content, Table S1) based on a modified version of Lewis food (Lewis, 1960, Table S1). These diets are a subset of ten diets used in an earlier study (Savola et al. 2020).

### LARVAL EXPERIMENTAL METHODS

*D. melanogaster* experimental individuals were from an outcross DGRP population established from 100 pairwise crosses of 113 lines from the *Drosophila* genetic reference panel (DGRP) (Mackay et al. 2012) (see methods and supplementary material in Savola *et al.* 2020). From the 35^th^ generation, we pipetted 5 μl of egg solution into each larval diet vial (following Clancy and Kennington, 2001) to establish density controlled groups of eggs (on average 50 (± 19) eggs). The mean volume of food in vials was 7.66 (± 0.58) ml. 30 vials of each diet were prepared in this way, and an additional 21 of the lowest P:C diet to ensure enough adults for adult collection. Due to very low egg counts in a small number of vials, a further 5-10 μl was added to these vials (17/171 vials, approximately 10%). For each vial, eggs were counted twice under a microscope to get an average egg count. Starting from experimental day 1 (one day after adding eggs to diets), vials were inspected daily for adult eclosion and the total number of adults eclosed per vial was recorded. The number of pupae per vial was counted once all adults had eclosed, based on pupal cases and undeveloped pupae.

### ADULT COLLECTION

Eclosion began on experimental day 8, from which point onwards adults were counted and removed from vials twice a day, in the morning and evening. Pilot data suggested different development times on the different diets, and therefore for each diet adults were collected primarily across three days per diet, starting one day after adult eclosion began (Figure S1). In this way, adult females were collected across a total of six days across all diet treatments to create six blocks. Some diets with quicker development times required some adults to be collected on the fourth day to achieve sufficient sample sizes (Figure S1 and Table S2; sample sizes = 18 to 40 adult flies per larval diet and stress treatment).

After adult collection, all flies were placed singly in vials containing standard Lewis medium in our laboratory, corresponding to the 1:6 P:C diet (Table S1, 14% protein diet), and were maintained on this diet for their remaining lifespan. Trays were rotated in the incubators daily to minimise microclimate effects. The day following adult collection, each female was provided with an age-matched male from the same outcross DGPR population. The female was left with the male for 24 hours to allow mating, following which the male was removed.

### STRESS TREATMENTS

On the seventh day post-eclosion for each block, female flies were exposed to one of three stress treatments: unstressed control, injury, and infection with a bacterial pathogen *Pseudomonas entomophila*. Treatments for each block were done at the same time each day (around 14:00) to minimise time-of-day effects on immunity (e.g. Lee and Edery, 2008). After stress treatments, the fly was placed into a new vial containing modified Lewis medium (Lewis, 1960, see P:C 1:6 diet in Table S1). The stress treatments were applied following Savola *et al.* (2020, modified from Dieppois *et al.* 2015; Troha and Buchon, 2019), with the overnight *P. entomophila* bacterial solution re-suspended in 30 ml Luria-Bertani (LB) medium in the morning and left to grow for three hours prior to dilution from a known OD value to correspond to an OD value of 0.001. This level was chosen from a pilot study (Figure S2) and is slightly lower than a previous experiment in adults (Savola et al. 2020), as the diluted OD of 0.005 had much lower survival compared to the previous experiment. Each block’s bacterial culture was established from a set of isogenic bacterial cultures grown overnight in LB medium, aliquoted in 20-25% glycerol 200 μl quantities and stored at −80°C. A subset of flies from each block of infections were plated on *Pseudomonas* isolating agar to confirm the infection treatments were successful (following Gupta *et al.* 2017, see supplementary information).

### ADULT TRAIT MEASUREMENTS

The number of eggs a female laid was counted from day 2 onwards for each block. For the first 14 days, eggs were counted daily and females were tipped into new vials. Subsequently, egg counts were performed every second day and stopped on day 98 for logistical reasons. This is an accurate proxy for lifetime egg production, as females in our previous experiment with adult P:C diet manipulation laid on average 99.37% (± 2.34%) of their lifetime eggs by day 98 (Savola et al. 2020). If a fly died on a day when eggs were not counted, an extra egg count was performed on the day of death. Survival was checked daily.

### STATISTICAL METHODS

The data were analysed using R software, version 3.5.2 (R Core Team 2014). All graphs were drawn using ggplot2 (Wickham 2016). All traits were analysed using generalised linear mixed models (GLMM). Models using a Poisson distribution were checked for zero inflation and overdispersion using the *DHARMa* package (Hartig 2019). In all models, even though we altered the P:C ratio of diets, diet was analysed as a continuous covariate as the percentage of protein in the diet (Table S1). To allow for non-linear effects, the quadratic term of protein percentage was also included. To avoid scaling errors, all continuous covariates were standardised to a mean of zero and a standard deviation of one. This was done separately for each test due to different sample sizes for different measures. Stress treatment was analysed as a categorical fixed effect. For all models with two-, or three-way interactions, for displaying summaries of LRT results of main effects or two-way interactions, parameter estimates and associated standard deviations are from separate models not including the associated two-, or three-way interactions. All model prediction plots were made with all random effects set to 1 and diets are shown as the percentage of protein in the diet.

We analysed survival to a number of developmental stages: egg-to-pupa, pupa-to-adult and egg-to-adult using linear models assuming a Gaussian distribution. For egg-to-pupa and egg-to-adult survival, we included the number of eggs in each vial as a covariate to control for differences in initial egg number on how many pupae or adults developed in each vial. This is essentially the same as modelling viability as percentages or as a proportion, which is often done in other larval diet studies (e.g. Andersen *et al.* 2010; Sentinella, Crean and Bonduriansky, 2013; Kutz, Sgrò and Mirth, 2019), however our method does not bound the data at 100% or 1. Similarly, for egg-to-pupa survival, we included the number of pupae in each vial as a covariate. All models were visually analysed for normality. Development time was analysed through GLMM with a Poisson error distribution using the lme4 R package (Bates et al. 2015) with the “bobyqa” optimiser with 100’000 iterations. We included vial as a random effect to account for any within vial effects, for example different larval densities and repeated measurements. For clarity of presentation, model predictions were made at either the average number of eggs or pupae per vial.

For adult survival, Kaplan-Meier survival curves were made using the survminer R package (Kassambara and Kosinski 2018) with diet as a factor. As our data did not conform to the assumptions of proportional hazards (global term of cox.zph function Chisq = 54.98, p = <0.001), we used an event history model following Moatt *et al.* (2019; Savola et al. 2020), implemented through a binomial GLMM in the lme4 R package (Bates et al. 2015) with the “bobyqa” optimiser with 100’000 iterations. This model is similar to the Cox proportional hazards model, however the results in this case estimate a per day mortality risk, which we will refer to as mortality. Individuals in the dataset were scored daily as 0 for alive and once as 1 for dead. The model included day as a random effect to account for differences in survival between days, and individual ID to account for multiple measures of an individual. To confirm the results of the survival model, we further analysed the data as lifespan using a GLMM in the package glmmTMB (Brooks et al. 2017) with a negative binomial distribution and block as a random effect to account for differences between infection days. Even though the data do not conform to the Cox proportional hazards assumptions, we checked for consistency with the result of the above analysis using a Cox proportional hazards model in the R Survival package (Therneau 2015).

For reproduction, various measures were analysed. As adult diet might have an increasing influence on the number of eggs produced as flies get older, for example due to compensatory feeding, we analysed measures of early-life egg production. We ran separate analyses on the number of eggs produced prior to stress treatments, seven days in total, and eggs produced in the seven days post stress treatments to test for early-life differences in reproduction and if these were affected by stress treatment. Only flies left alive on the last day of egg counts were included in these analyses. These data were analysed using a GLMM with a negative binomial distribution and including a zero-inflation term with the glmmTMB R package (Brooks et al. 2017). Lifetime egg production (to day 98) was analysed with an identical model to the other reproduction models with all flies included, with an additional model including lifespan as a predictor. Mean centered lifespan was included to account for selective disappearance and block was included as a random effect. To analyse daily egg production, all egg counts corresponding for a span of two days were divided by two and rounded down to the nearest integer to match earlier daily egg counts. Daily egg production was analysed using a GLMM with negative binomial distribution and including a zero-inflation term. As well as the fixed effects described above, age was included as a linear and non-linear term as well as the interactions between these and all other fixed effects. Individual ID and block were included as random effects.

## RESULTS

### EFFECTS OF LARVAL NUTRITION ON LARVAL TRAITS

P:C of the larval diet had a significant effect on how many individuals developed from eggs to adults, where larvae reared on higher P:C were more likely to develop to adults (Figure 1A & Figure S3A; Table S3A; Protein = 2.20 (± 0.55), F = 15.90, p = <0.001). With higher numbers of eggs in a vial, more adults developed (Table S3A; Average number of eggs = 0.72 (± 0.03), F = 762.19, p = <0.001). Separating this result into effects on larval and pupal viability, there was a significant effect of P:C on the numbers of eggs developing into pupae (Figure 1B & Figure S3B; Table S3A; Protein = 1.79 (± 0.52), F = 9.13, p = 0.003; Average number of eggs = 0.86 (± 0.03), F = 1165.5, p = <0.001). There was a marginally non-significant effect of P:C on the number of adults developing from pupae (Figure 1C & Figure S3C; Table S3C; Protein = 0.70 (± 0.45), F = 3.81, p = 0.052). As expected, with more pupae in a vial, more adults developed (Figure 1C & Figure S3C; Table S3C; Pupae = 0.82 (± 0.02), F = 1228.5, p = <0.001).

**Figure 1:**
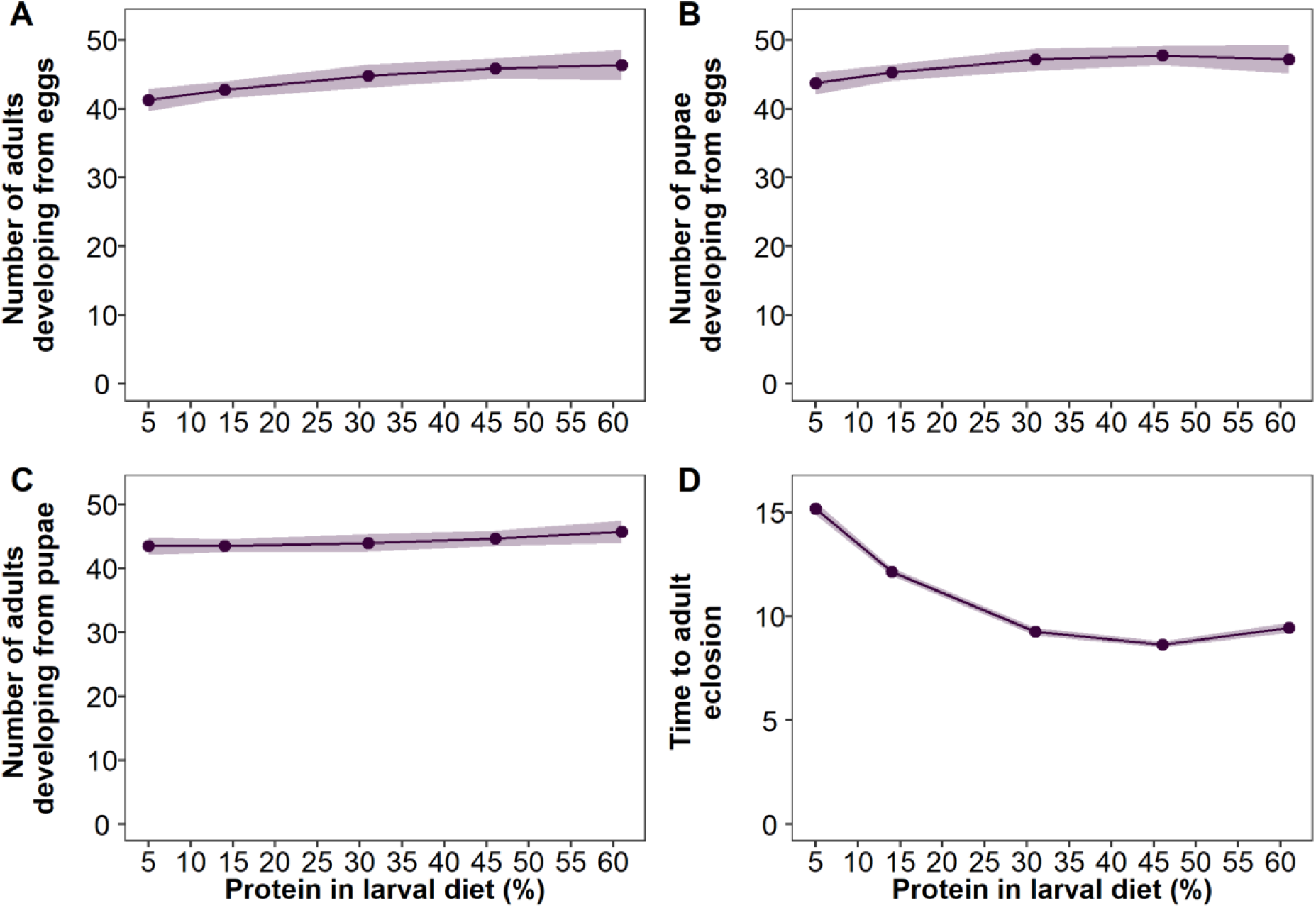
Model predictions of the effects of larval P:C (shown as the corresponding percentage of protein) on various larval traits: (A) the number of adults developing having controlled for the number of eggs laid (50 (± 19) eggs on average); (B) the number of pupae eclosing having controlled for the number of eggs laid (50 (± 19) eggs on average); (C) the number of adults developing having controlled for the number of pupae formed (46 (± 18) pupae on average); and (D) the average time taken for adult eclosion. All predictions are based on either vials starting with the overall mean number of eggs (50 (± 19) eggs) (A, B, D), or the overall mean number of pupae (46 (± 18) pupae) (C). Shaded areas are 95% confidence intervals. Protein and protein^2^ are mean centered to standard deviation of 1. See S3 for viability data as percentages.

P:C in the larval diet also had an effect on the development time to adulthood, with higher larval P:C resulting in shorter development time (Figure 1D & Figure S4, Table S3; Protein = −0.22 (± 0.01), Chi-squared = 191.75, p = <0.001). This relationship is quadratic, suggesting that intermediate P:C diets had a quicker development time in comparison to the high or low P:C diets, or that the rate of reduction in development time plateaued at the highest P:C diets (Figure 1D & Figure S4, Table S4; Protein^2^ = 0.16 (± 0.01), Chi-squared = 183.61, p = <0.001). Vials with higher average number of eggs had a slightly longer development time (Table S4; Average number of eggs = 0.03 (± 0.01), Chi-squared = 26.80, p = <0.001).

### EFFECTS OF LARVAL NUTRITION ON ADULT TRAITS AND SURVIVAL AFTER STRESS

P:C of larval diet had no effect on adult mortality regardless of stress treatment (Figure 2B & Figure S5; Table S5 & Table S6). Stress treatment had a significant effect on mortality, where infected flies had a higher risk of death (Figure 2B & Figure S5; Table S5; Treatment Chi-squared = 76.67, p = <0.001; Infection = 1.18 (0.13); Injury = 0.24 (± 0.17)). Analysing the survival data as lifespan, the same patterns were observed, where stress treatment had a significant effect on lifespan, with infected flies having shorter lifespans (Figure S6 & Figure S7; Table S7 & Table S8; Treatment Chi-squared = 99.28, p = <0.001; Infection = −1.00 (± 0.10); Injury = −0.10 (± 0.10)). The results of a Cox proportional hazards model show the same patterns (Figure S8; Table S9).

**Figure 2:**
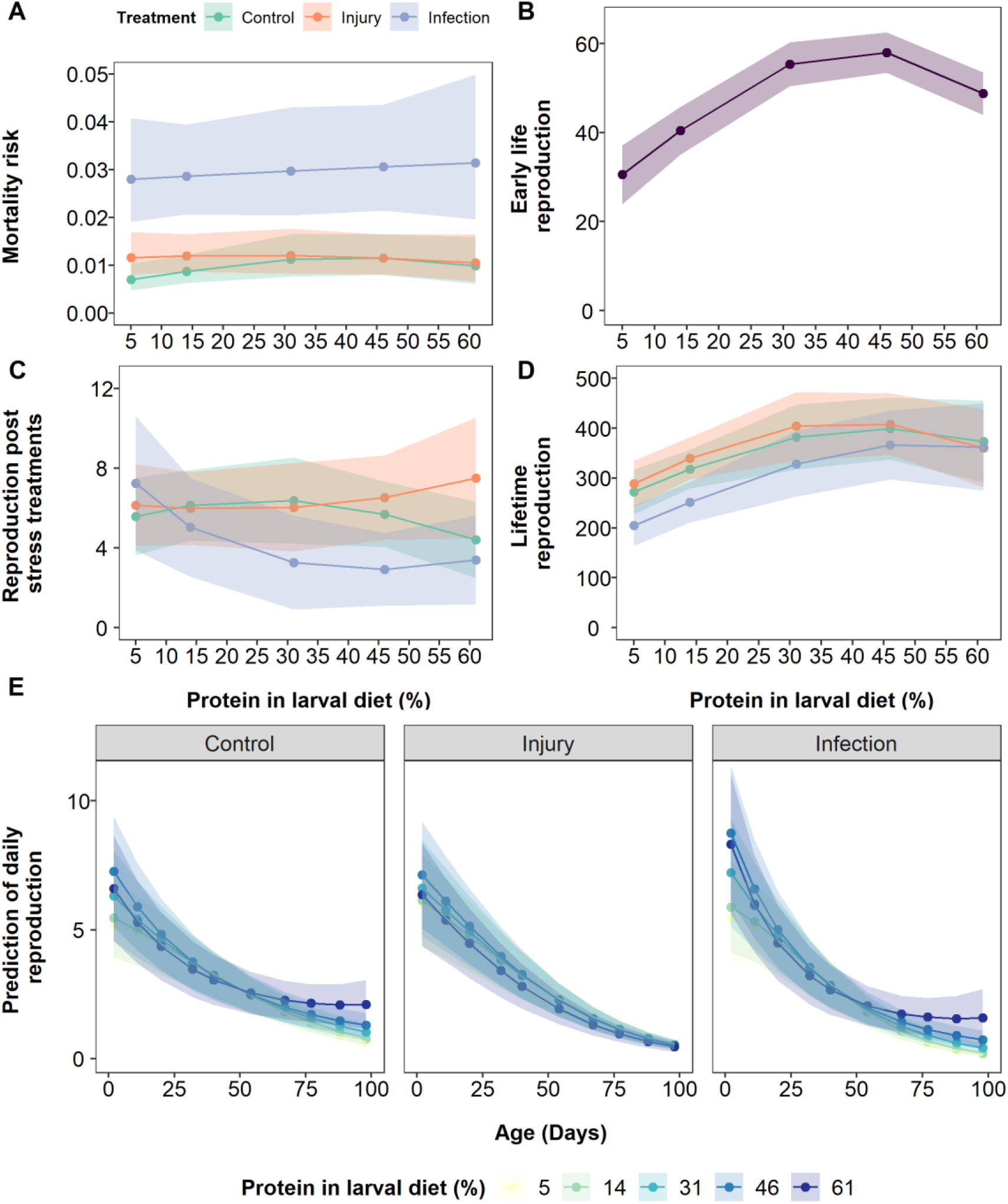
Model predictions of the effects of larval diet P:C (shown as the corresponding percentage of protein) and adult stress treatment on various adult life-history traits: (A) per day mortality risk; (B) egg production in the first 7 days of adulthood (prior to stress treatments); (C) egg production across the 7 days after stress treatments; (D) lifetime egg production (up to day 98); (E) reproductive ageing in terms of daily egg production. Adult flies were infected with a bacterial pathogen (blue data points and lines in A, C, D), injured by a pinprick (orange data points and lines in A, C, D) or with no treatment (green data points and lines in A, C, D). Lifespan is accounted in the model to account for selective disappearance in (D). Shaded areas are 95% confidence intervals. Protein and protein^2^ (A-E), and age and age^2^ (E) are mean centered to standard deviation of 1.

Larval P:C had a significant effect on early-life egg production prior to stress treatments, where increasing larval P:C increased early-life egg production which then levelled off at very high P:C diets (Figure 2B & Figure S11; Table S14; Protein = −0.29 (± 0.06), chi-squared = 3.71, p = 0.054; Protein^2^ = −0.20 (± 0.04), p = <0.001). There were no significant effects of larval diet or stress treatments on egg production in the seven days following stress treatments (Figure 2C & Figure S12; Table S15 & Table S16).

The effect of larval P:C on lifetime egg production was similar to the effect on early-life reproduction, where increasing P:C in the larval diet increased lifetime egg production (Figure 2D & Figure S9; Table S10; Protein = 0.11 (± 0.05), Chi-squared = 5.73, p = 0.02). The effect of P:C was non-linear (Figure 2D; Table S10; Protein^2^ = −0.11 (± 0.04), Chi-squared = 8.01, p = 0.005), with egg production reaching a peak at intermediate P:C and not increasing further at higher P:C. Stress treatment had a significant effect on lifetime egg production, where infected flies produced fewer eggs (Figure 2D; Figure S9; Table S10; Treatment chi-squared = 8.81, p = 0.01; Infection = −0.18, (± 0.08); Injury = 0.04 (± 0.07)). Flies produced more eggs with longer lifespan (Table S11, Lifespan = 0.69 (± 0.04), Chi-squared = 275.77, p = <0.001). There was no significant interaction between larval P:C and stress treatment (Table S11). A model not accounting for lifespan showed the same pattern with stress treatments having a significant effect on egg production, where infected individuals produced fewer eggs (Figure S10; Table S12 & Table S13; Stress treatment chi-squared = 80.68, p = <0.001; Infection = −0.82 (± 0.09); Injury = −0.04 (± 0.09)). Increasing larval P:C resulted in higher lifetime egg production, however this pattern was not quadratic (Figure S10; Table S12 & Table S13; Protein = 0.12 (± 0.04), Chi-squared = 5.91, p = 0.02; Protein^2^ = 0.07 (± 0.05), Chi squared = 2.10, p = 0.15). Even though the models differ slightly, the P:C patterns are broadly similar for both models, where egg production plateaus at the highest P:C level, as also seen in the raw data (Figure 2D, S9 & S10).

In general, flies across all larval diets and stress treatments showed similar patterns in egg laying over their lifespan (Figure 2E & Figure S13). Egg production was highest early in life and then declined (Figure 2E & Figure S13). At mean centered P:C and lifespan, there was a slowing in the rate of decline of egg laying with age, such that the rate of decline is high when young and slows as flies get older (Table S17 & Table S19; Age = −0.60 (± 0.03), chi-squared = 2175.5, p = <0.001; Age^2^ = −0.03 (± 0.02), chi-squared = 26.14, p = <0.001). As with lifetime egg production, there was a significant non-linear effect of P:C, with intermediate P:C diets resulting in higher daily egg production in the control stress treatment at mean age and lifespan (Figure S13; Table S17 & Table S19; Protein^2^ = −0.05 (± 0.05), chi-squared = 12.26, p = 0.0005). Lifespan had no effect on daily egg production (Table S19, Lifespan = 0.02 (± 0.02), p = 0.16).

The pattern of ageing in reproduction was broadly similar across larval diets and adult stress treatments (Figure 2E & Figure S13). However, there were some significant two-, and three-way interactions, which indicate small differences in the pattern of reproductive ageing across diets and stress treatments. There was a significant two-way interaction between P:C and the linear and quadratic effect of age, suggesting that with higher P:C, ageing in egg production was quicker and the linear effect of age was highest at intermediate P:C diets (Figure S13; Table S18 & Table S19; Protein:Age = −0.10 (± 0.02), chi-squared = 30.48, p = <0.001; Protein:Age^2^ = 0.10 (± 0.02), chi-squared = 23.94, p = <0.001). There was also a significant two-way interaction between age and the quadratic effect of P:C, suggesting that the rate of ageing was highest at intermediate P:C (Figure S13; Table S18 & Table S19; Protein^2^:Age = 0.12 (± 0.02), chi-squared = 22.62, p = <0.001). Stress treatments had significant effects on these two-way interactions (Figure 2E & Figure S13; Table S19, Treatment:Protein:Age^2^ chi-squared = 14.41, p = 0.001, Treatment:Protein^2^:Age chi-squared = 11.21, p = 0.004), where in injured flies these terms were smaller compared to the control individuals (Table S19; Injury:Protein:Age^2^ = −0.08 (± 0.03); Injury:Protein^2^:Age = −0.10 (± 0.03)). This suggests that there was less of an effect of P:C on aging in the injured flies. Stress treatment also had a significant effect on ageing, where infected flies had a more negative decline in egg laying with age than control individuals (Figure 2E & Figure S13; Table S19; Treatment:Age chi-squared = 18.05, p = <0.001; Infection:Age = −0.25 (± 0.06)).

## DISCUSSION

The main objective of our study was to test whether larval diets ranging in P:C content affected adult survival post-infection. We predicted that adults that developed on lower P:C diets would have worse survival post-infection, due to the demonstrated importance of dietary protein for the response to infection (e.g. Lee *et al.* 2006; Povey *et al.* 2009; Cotter *et al.* 2019; Savola *et al.* 2020). However, our results provide no evidence for an effect of larval P:C diet on adult lifespan, regardless of stress treatment. Similarly, although intermediate P:C in the larval diet increased lifetime and early-life reproduction, there were no interactions between larval diet and stress treatment on reproduction, except for a smaller effect of P:C on the senescent decline in egg production for injured flies. Infection did however have effects on many life-history traits, specifically reducing lifespan and lifetime egg production, and increasing the rate of senescence in egg laying. These results suggest that, in this study, although P:C in larval diet affected larval and adult life-history traits, and exposure to infection in adulthood affected adult life-history traits, these effects did not interact very strongly.

Previous studies have suggested links between larval diet and the ability of adults to cope with environmental stress such as infection. For example, lower P:C nymphal diet with matching adult diet decreased adult survival post-infection measured for five days in female *Gryllus texensis* crickets (Kelly and Tawes 2013). Similarly, adult *Anopheles gambiae* mosquitoes raised as larvae on increasing CR had lower melanising ability (Suwanchaichinda and Paskewitz 1998). Studies measuring components of the immune system without a direct stressor have also shown that adults raised on non-optimal diets had lower adult immune function, with short term starvation (De Block and Stoks 2008) or caloric restriction (Rolff et al. 2004). Similarly, adult *D. melanogaster* raised on higher P:C diets had higher levels of *Diptericin A* and *Metchnikowin* AMP transcription (Fellous and Lazzaro 2010). This is particularly relevant to our study, as AMPs are important in bacterial defence (reviewed in Zhang and Gallo, 2016) and in *D. melanogaster* AMPs including *Metchnikowin* and *Diptericin* are upregulated with *P. entomophila* infection (Liehl et al. 2006; Chakrabarti et al. 2012). However, in this previous study, only the amount of yeast in diet was altered without reducing carbohydrates thus altering both calorie and P:C content. Consequently, increased calories may be driving these effects and not the increase in P:C. As previous studies have all manipulated calories, perhaps the caloric value of juvenile diet affects adult immunity and not the macronutrient content. By only manipulating the P:C content of larval diets, our results suggest that macronutrient ratio does not affect adult survival post-infection.

Outside of differences in the type of diet manipulation, another factor that could explain our contrasting findings to previous research is the type of stress experienced. For example, *D. melanogaster* flies raised on higher P:C had a longer chill coma recovery time (Andersen et al. 2010) and worse starvation resistance (Davies et al. 2018), however better heat coma and desiccation resistance (Andersen et al. 2010). It has been suggested that these larval diet effects are a result of effects on general body condition or specific tissues (Fellous and Lazzaro 2010), the production of heat shock proteins (Andersen et al. 2010), and more directly, lipid storage through eating a diet richer in carbohydrates as larvae (e.g. Roeder and Behmer, 2014; Kim, Jang and Lee, 2020). Our results suggest this does not seem true for immune responses, even though the fat body is an organ linked to both larval feeding and immune responses (reviewed in Arrese and Soulages, 2010). Larval feeding may therefore differentially affect adult stress response where certain diets are better for specific environmental stressors.

Another consideration is the timing of the stressor, as previous studies have applied stressors closer to eclosion. Our seven day lag post-eclosion could allow enough time for compensatory mechanisms (reviewed in Nestel *et al.* 2016), such as compensatory feeding (Raubenheimer and Simpson 1993) to mask any effects of larval diet. For example, adults that developed on low P:C diets could have eaten enough protein to survive injury and infection to a similar level to adults that developed on higher P:C. Two studies that altered both larval and adult diets found that adult environment was the main determinant of life-history traits (Davies et al. 2018; Duxbury and Chapman 2020), however, in one there were small and complex differences in female lifetime reproduction between larval and adult diet combinations (Duxbury and Chapman 2020). It would therefore be interesting to repeat our experiment and expose adults to injury and infection stress immediately upon eclosion.

There was no effect of larval diet on survival or lifespan, as seen in other studies (Tu and Tatar 2003; Houslay et al. 2015; Davies et al. 2018). In adults, altering P:C affects lifespan, with lifespan maximised on either intermediate (e.g. Lee, 2015; Kim, Jang and Lee, 2020; Savola *et al.* 2020) or low P:C diets (e.g. Lee *et al.* 2008; Maklakov *et al.* 2008; Jensen *et al.* 2015). A small number of studies have suggested an effect of larval diet P:C on adult lifespan, but results have been inconsistent (lifespan maximised on high (Duxbury and Chapman 2020), intermediate (Runagall-McNaull et al. 2015; Kim et al. 2019), and low (Economos 1984; Stefana et al. 2017) P:C diets). A more consistent role has been suggested for calories, with adult lifespan decreasing with larval calorie restriction (May et al. 2015; Adler et al. 2016; Hooper et al. 2017; Krittika et al. 2019). Given a meta-analysis found no overall effect of early-life diet on lifespan (English and Uller 2016), such contrasting findings suggest no clear effect of larval diet on adult lifespan and instead suggest that lifespan is more determined by adult diet.

Lifetime and early-life reproduction increased with increasing larval P:C and then declined slightly at the highest P:C. Similarly, ovariole number has been show to peak at intermediate larval P:C in *Zaprionus indianus* (Matavelli et al. 2015) and in *Drosophila melanogaster* (Rodrigues et al. 2015). Many larval studies lack this decline at the highest P:C diets (e.g. Tu and Tatar, 2003; Andersen *et al.* 2010; Silva-Soares *et al.* 2017; Duxbury and Chapman, 2019; Kim *et al.* 2019). Protein is often the limiting nutrient in egg production (reviewed in Wheeler, 1996; Boggs, 2009), but can have a toxic effect when consumed at very high levels (reviewed in Simpson and Raubenheimer, 2009), which may explain the plateauing at very high P:C due to larvae of worse condition developing to adults. Due to the highest P:C ratio also including the lowest carbohydrate content, this effect may also be due to a limiting effect of carbohydrates on development. As nutrients for egg production can be acquired from adult feeding and nutrient requirements can differ between species (reviewed in Wheeler, 1996), this limit may not always appear. These P:C effects could also arise through the general increase in body condition with higher P:C in larval diet (Runagall-McNaull et al. 2015). Overall, there appears to be an increase in adult reproduction with increasing P:C in the larval diet, but this effect may plateau at very high P:C levels.

Infection reduced lifetime egg production, a typical response in insects (reviewed in Schwenke et al. 2016), however there was no interaction between larval P:C and reproduction post-infection. When accounting for the overall shorter lifespan of infected flies, their reproduction was comparable to the injured or unstressed flies. In the week after stress treatments, all treatment groups produced the same number of eggs. This suggests that with infection, individuals were able to produce more eggs earlier in life but then egg numbers declined. This was reflected in the reproductive ageing results, as infection increased the rate of senescence in egg laying. This could be evidence of terminal investment (Clutton-Brock 1984), specifically fecundity compensation, as flies shifted their egg production earlier as a response to infection, as is a common outcome after infection (reviewed in Kutzer and Armitage, 2016).

In general, the patterns of reproductive ageing were quite similar across treatments, but there were some interactions between P:C in the larval diet and stress treatment. Overall, at intermediate P:C, ageing in egg production was quickest. Similar results in reproductive ageing have been found in studies altering adult P:C (e.g. Jensen *et al.* 2015; Savola *et al.* 2020). These results are also similar to previous studies focusing on ageing in egg laying where adults raised on higher P:C and/or calories as larvae appear to have quicker ageing in egg laying (Tu and Tatar, 2003; Hooper *et al.* 2017, but see May, Doroszuk and Zwaan, 2015). As these studies manipulated calories, here we show there are minor changes in ageing patterns also with P:C manipulation of the larval diet.

For the larval traits, we predicted that larvae would have more successful and quicker development on higher P:C diets (Britton and Edgar 1998; Colombani et al. 2003; Chang 2004; Mirth and Riddiford 2007). Development time was quickest on intermediate P:C, and egg-to-pupae and egg-to-adult viability were higher on higher P:C, as seen in previous studies (e.g. Andersen et al. 2010; Silva-Soares et al. 2017; Kutz et al. 2019, but see Houslay *et al.* 2015; Davies *et al.* 2018; Gray, Simpson and Polak, 2018). However, despite being statistically different, the effect of diet on egg-to-pupae and egg-to-adult viability was small. There was no effect of P:C on pupae-to-adult viability, suggesting that all of the diets used in our study allowed larvae to pupate successfully. Studies often do not report pupae-to-adult viability, however more extreme diets could affect this trait as different sources of carbohydrates have been shown to affect pupae-to-adult viability (Nash and Chapman 2014). On the highest P:C diet, development time was slightly slower, which is also a common finding (e.g. Lee et al. 2012; Rodrigues et al. 2015; Kutz et al. 2019), again potentially due to toxic effects of high P:C diets (reviewed in Simpson and Raubenheimer, 2009) or due to a limitation of carbohydrates for development. Vials with more eggs took longer to develop, most likely due to larval density effects (e.g. Ludewig *et al.* 2017; Klepsatel, Procházka and Gáliková, 2018; Henry, Tarapacki and Colinet, 2020; Than, Ponton and Morimoto, 2020). Our work adds to growing evidence of the importance of macronutrients in larval diet for larval development and suggest that, with some exceptions, intermediate P:C diets are better for key larval traits.

## CONCLUSIONS AND FUTURE WORK

The results of this study suggest that larval dietary P:C has no effect on adult survival with or without stress treatment, and thus larval P:C does not alter the long-term consequences of injury or infection on survival. Intermediate P:C larval diets were optimal for many traits pre-, and post-metamorphosis. Individuals were the quickest to develop into adults on intermediate larval P:C, and subsequent adults produced the most early-life and lifetime eggs. Larvae were more likely to develop into adults on higher P:C. Therefore, our results add to the growing evidence that larval diet affects adult life-history traits, but the long-term consequences of infection and injury are not altered. To understand the effects of larval diet on the ability of adults to respond to infection further, we suggest experiments exposing adults to infection immediately after eclosion to avoid potential for compensatory feeding. Furthermore, using a fully factorial experiment combining variation in larval and adult diet could help to disentangle the differences between larval and adult feeding on stress responses such as infection and injury.

## ACKNOWLEDGEMENTS

This work was supported by the Biotechnology and Biological Sciences Research Council (BBSRC) grant number BB/M010996/1. We thank Joshua Moatt for helpful advice and comments on the data analysis and write-up, Katy Monteith for laboratory training and practical advice as well as help with data collection at the end of the experiment along with Paul McNeil and Paulina Mika, and Anu Halonen for data on confirmation of infection from one day of missing data.

## AUTHORS’S CONTRIBUTIONS

ES, PV and CW designed the experiment. ES did the study and analysed the data with help from CW. ES wrote the paper with help from CW and PV.

