## Supplementary information for "Larval diet affects adult reproduction but not survival regardless of injury and infection stress in *Drosophila melanogaster*"

SUPPLEMENTARY METHODS:

**Table S1:** Information on the five diets as a subset of ten diets used in (Savola et al., 2020). The standard Lewis food and associated P:C ratio is in bold (Lewis, 1960). One of the main differences to the original Lewis food recipe is the replacement of dextrose and sucrose with brown sugar in our diets (Lewis, 1960). The P:C ratios (rounded to the nearest whole number) incorporate the protein and carbohydrate contributed by maize. Yeast and sugar are roughly isocaloric, so P:C ratios can be altered without altering the energy content of the diet by replacing yeast with sugar (Mair et al., 2005).

| P:C ratio | Protein in diet (%) | Yeast (g) | Sugar (g) | Maize (g) | | | Agar (g) | Nipagin (ml) | $dH_2O$ (l) |
|---|---|---|---|---|---|---|---|---|---|
| | | | | Total | Carbohydrate | Protein | | | |
| 1:16 | 5 | 21.3 | 653.7 | 415 | 290.5 | 37.8 | 41.2 | 90 | 6 |
| **1:6** | **14** | **112.5** | **562.5** | **415** | **290.5** | **37.8** | **41.2** | **90** | **6** |
| 1:2 | 31 | 296.7 | 378.3 | 415 | 290.5 | 37.8 | 41.2 | 90 | 6 |
| 1:1 | 46 | 463.9 | 211.1 | 415 | 290.5 | 37.8 | 41.2 | 90 | 6 |
| 2:1 | 61 | 631.1 | 43.9 | 415 | 290.5 | 37.8 | 41.2 | 90 | 6 |

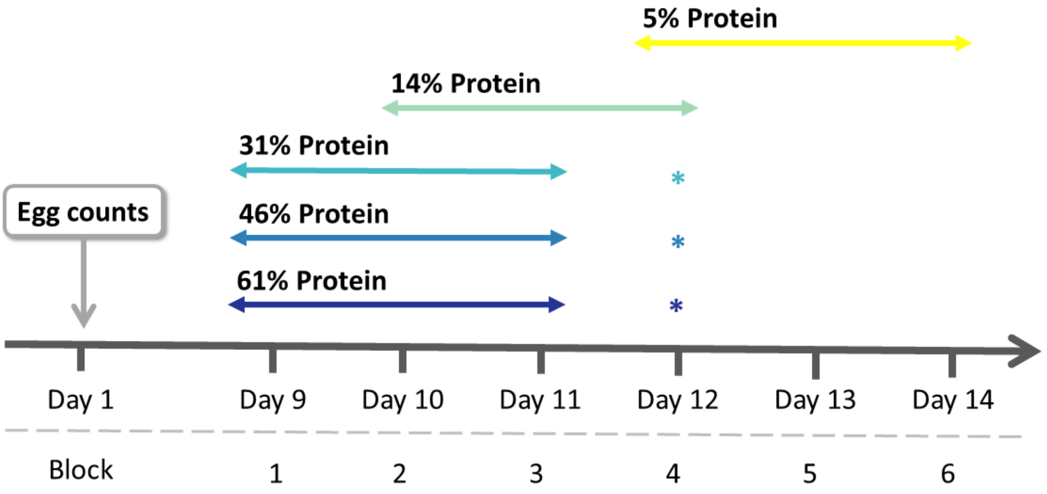


**Figure S1:** Schematic for adult collection across days to create 6 blocks of females. Stars (*) indicate if only a few addition adults were collected on this day to reach sample size per diet (see methods).

**Table S2**: Total sample size per diet and treatment of flies collected across three to four days after eclosion started.

| Protein in diet (%) | P:C ratio | Stress treatment | | |
|---|---|---|---|---|
| | | Control | Injury | Infection |
| 5 | 1:16 | 35 | 40 | 35 |
| 14 | 1:6 | 32 | 36 | 30 |
| 31 | 1:2 | 25 | 24 | 25 |
| 46 | 1:1 | 19 | 19 | 18 |
| 61 | 2:1 | 22 | 24 | 23 |

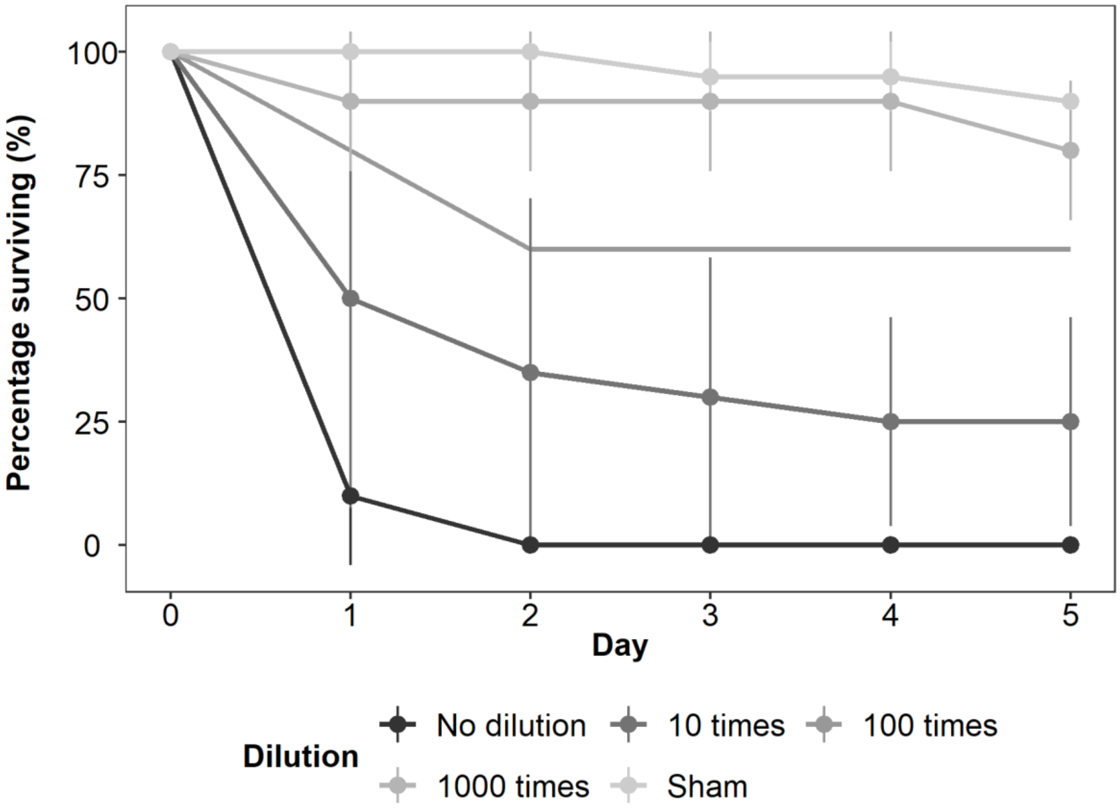


**Figure S2:** Dilution series for *Pseudomonas entomophila* bacterial solution from the same stock as used in infections. 10 females per vial were infected with the specified solution (no dilution to 1000 times dilution) or with no pathogen ("Sham"). Results show mean survival of two replicates of ten flies and the vertical lines indicate standard deviation, except for the 100 times dilution, which only has one replicate.

**Bacterial growth (CFU) measurements:**

24 hours post-infection two replicate groups of three flies from the infected, sham and control groups were plated (following Gupta et al., 2017). Across infection blocks, colonies grew on the plates confirming successful infections, except for the first block where initially only one fly per sample was used for the plating. Infected flies from the first block showed similar levels of mortality to flies from other blocks, suggesting they were indeed infected and that use of only a single fly resulted in bacterial levels that were below a detection threshold in the assay. Due to logistical reasons, the last block of infections was plated 48 hours post-infection, however another group of infected flies from the same overnight bacterial culture showed growth (Halonen, data not shown).

SUPPLEMENTARY RESULTS:

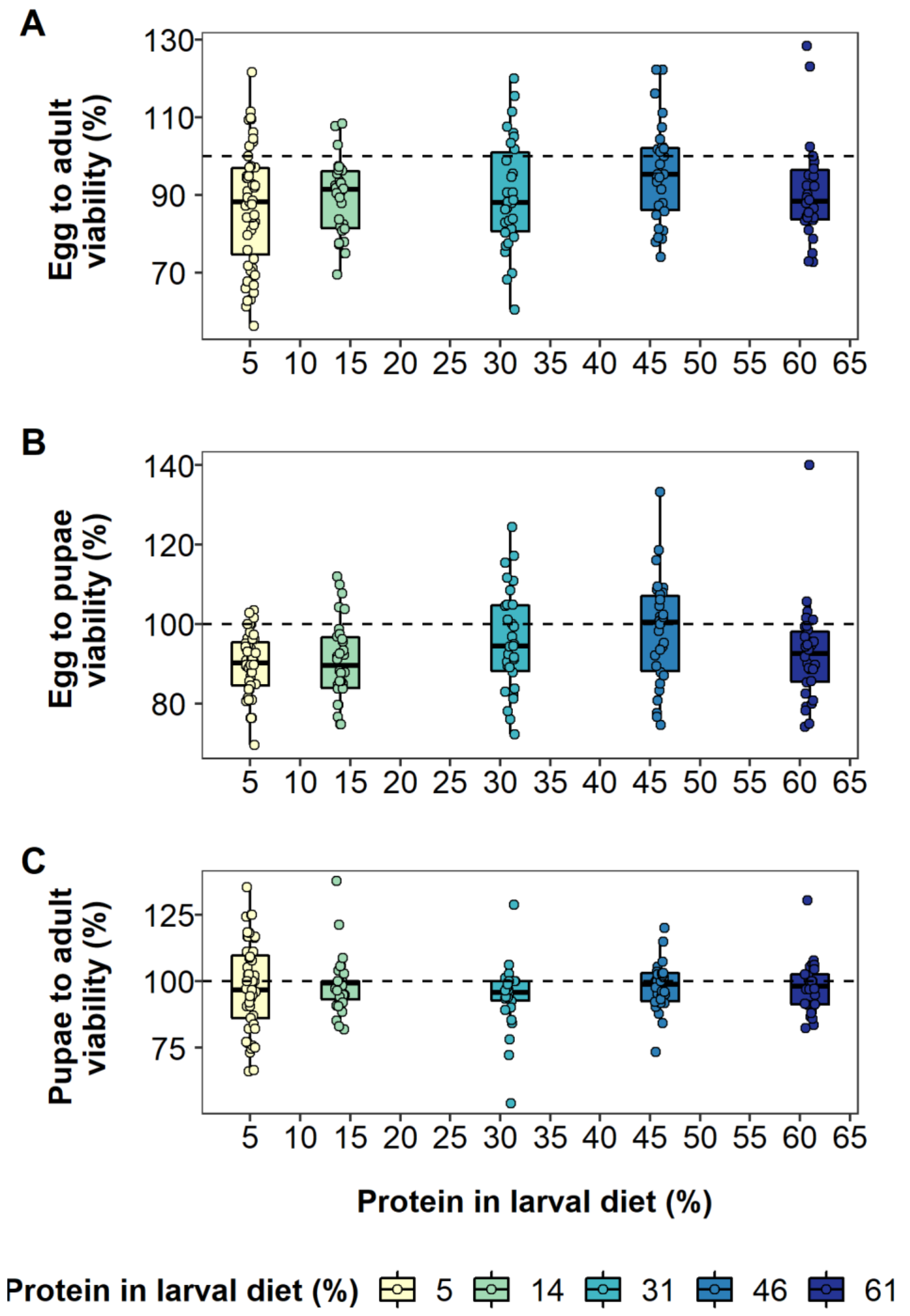


**Figure S3:** Effects of protein in larval diet on the percentage of eggs developing to adults (A), eggs developing to pupae (B) and pupae developing to adults (C). Values are over 100% due to inaccuracies in egg and pupal counts. The lines in the box plots indicate median values (50% quantile), boxes are the interquartile range (25% to 75% quantiles) and whiskers are minimum or maximum quartiles (25% - 1.5 x interquartile range, 75% + 1.5 x interquartile range).

**Table S3:** Model summary of a Gaussian linear model of the effects of protein in larval diet and the number of eggs laid in the vial (averaged over two counts, see methods) on the number of adults developing per vial (A); the number of eggs laid in the vial on the number of pupae developing per vial (B); and the number of pupae in the vial on the number of adults developing per vial (C). Protein and protein$^2$ are mean centered to standard deviation of 1. Significant results below significance level $\alpha = 0.05$ are bolded.

| (A) Number of adults developing from eggs: | | | | | |
|---|---|---|---|---|---|
| | Estimate | Standard error | Df | F | Pr (>F) |
| Intercept | 8.27 | 1.54 | | | |
| **Average number of eggs** | **0.72** | **0.03** | **1** | **762.19** | **<0.001** |
| **Protein** | **2.20** | **0.55** | **1** | **15.90** | **<0.001** |
| Protein$^2$ | -0.66 | 0.69 | 1 | 0.91 | 0.34 |
| (B) Number of pupae developing from eggs: | | | | | |
| | Estimate | Standard error | Df | F | Pr (>F) |
| Intercept | 3.99 | 1.47 | | | |
| **Average number of eggs** | **0.86** | **0.03** | **1** | **1165.5** | **<0.001** |
| **Protein** | **1.79** | **0.52** | **1** | **9.13** | **0.003** |
| Protein$^2$ | -1.05 | 0.66 | 1 | 2.57 | 0.11 |
| (C) Number of adults developing from pupae: | | | | | |
| | Estimate | Standard error | Df | F | Pr (>F) |
| Intercept | 6.15 | 1.27 | | | |
| **Pupae** | **0.82** | **0.02** | **1** | **1228.5** | **<0.001** |
| Protein | 0.70 | 0.45 | 1 | 3.81 | 0.052 |
| Protein$^2$ | 0.31 | 0.56 | 1 | 0.31 | 0.58 |

**Table S4:** Model summary of a Poisson model of the effects of protein in larval diet and the average number of eggs laid in the vial on the number of days until adult eclosion. Vial ID was fitted as a random effect. Protein, protein$^2$ and average egg counts are mean centered to standard deviation of 1. Significant results below significance level $\alpha = 0.05$ are bolded.

| | Estimate | Standard error | Z value | Df | Chisq | Pr (>Chisq) |
|---|---|---|---|---|---|---|
| Intercept | 2.23 | 0.01 | 206.87 | | | |
| **Protein** | **-0.22** | **0.01** | **-35.32** | **1** | **191.75** | **<0.001** |
| **Protein$^2$** | **0.16** | **0.01** | **18.33** | **1** | **183.61** | **<0.001** |
| **Average number of eggs** | **0.03** | **0.01** | **5.39** | **1** | **26.80** | **<0.001** |

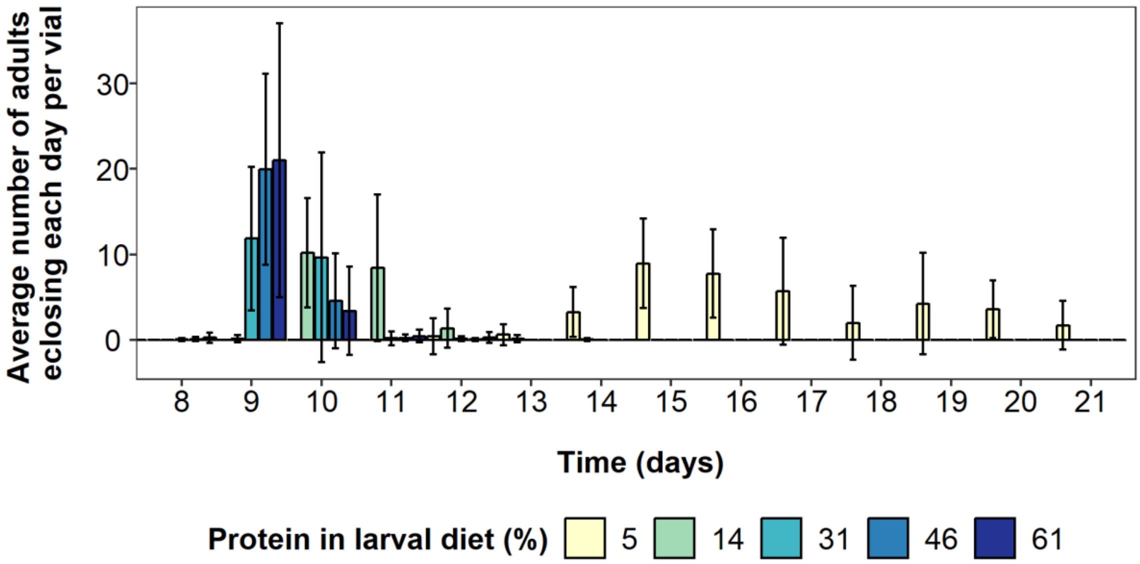


**Figure S4:** Effects of protein in larval diet on the average number of adult flies eclosing each day after egg production. No adults eclosed prior to day 8, so these days are not shown. Error bars are standard deviations.

**Table S5:** Summary of main effects parameter estimates and associated LRT test values for a binomial model of the effects of protein in larval diet and stress treatments on mortality risk per day. The values are from models not including interactions with the specific main effect. Chi-squared and associated p-values are from LRT tests comparing a model with no interactions associated with the main effect to a model with no main effect. Protein and protein$^2$ are mean centered to standard deviation of 1. Significant results below significance level $\alpha$ = 0.05 are bolded.

| | Estimate | Standard error | Z value | Df | Chisq | Pr (>Chisq) |
|---|---|---|---|---|---|---|
| **Injury treatment** | **0.24** | **0.17** | **-27.26** | **2** | **76.67** | **<0.001** |
| **Infection treatment** | **1.18** | **0.13** | **1.87** | | | |
| Protein | 0.05 | 0.05 | 0.98 | 1 | 0.88 | 0.35 |
| Protein$^2$ | -0.07 | 0.07 | -1.01 | 1 | 0.98 | 0.32 |

**Table S6:** Model summary of a binomial model of the effects of protein in larval diet and stress treatments on mortality risk per day. Protein and protein$^2$ are mean centered to standard deviation of 1.

| | Estimate | Standard error | Z value | Df | Chisq | Pr (>Chisq) |
|---|---|---|---|---|---|---|
| Intercept | -4.52 | 0.19 | -23.34 | | | |
| Injury treatment | 0.12 | 0.21 | 0.57 | | | |
| Infection treatment | 1.02 | 0.21 | 4.94 | | | |
| Protein | 0.23 | 0.11 | 2.13 | | | |
| Protein$^2$ | -0.17 | 0.12 | -1.40 | | | |
| Injury:Protein | -0.24 | 0.15 | -1.60 | 2 | 2.52 | 0.28 |
| Infection:Protein | -0.18 | 0.15 | -1.22 | | | |
| Injury:Protein$^2$ | 0.12 | 0.17 | 0.74 | 2 | 0.97 | 0.62 |
| Infection:Protein$^2$ | 0.17 | 0.17 | 1.00 | | | |

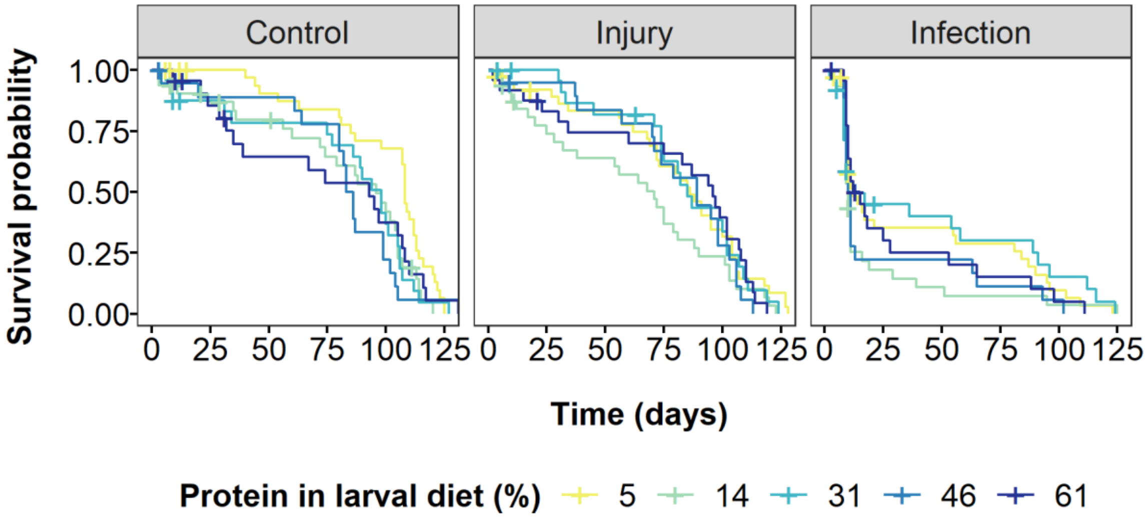


**Figure S5:** Effects of protein in larval diet on survival of adult flies infected with a bacterial pathogen ("Infection"), injured by a pinprick ("Injury") or with no treatment ("Control"). Survival is shown as Kaplan-Meier curves for each stress and diet treatment groups. Plus signs (+) indicate censored data points.

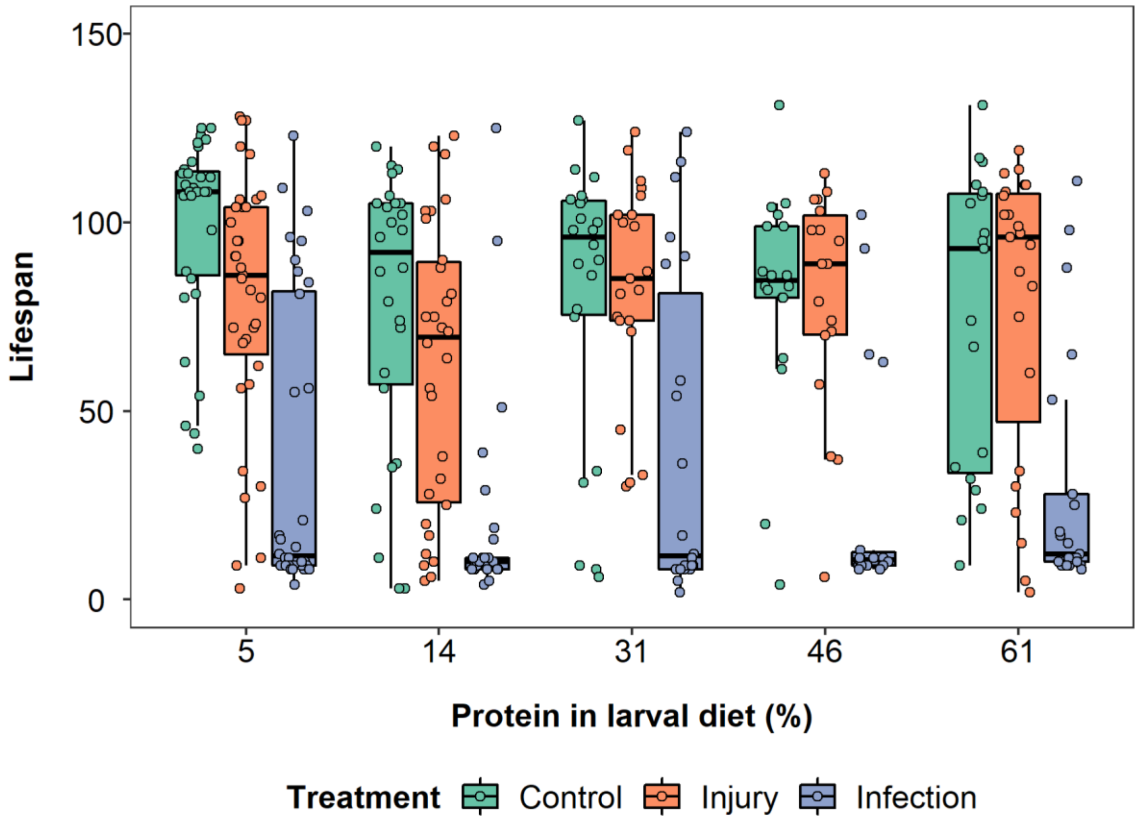


**Figure S6:** Effects of protein in larval diet on the lifespan of flies infected with a bacterial pathogen (blue bars and data points), injured by a pinprick (orange bars and data points) or with no treatment (green bars and data points). The lines in the box plots indicates median values (50% quantile), boxes are the interquartile range (25% to 75% quantiles) and whiskers are minimum or maximum quartiles (25% - 1.5 x interquartile range, 75% + 1.5 x interquartile range).

**Table S7:** Summary of main effects parameter estimates and associated LRT test values for a negative binomial model of the effects of protein in larval diet and stress treatments on lifespan of adult flies. The values are from models not including interactions with the specific main effect. Chi-squared and associated p-values are from LRT tests comparing a model with no interactions associated with the main effect to a model with no main effect. Protein and protein$^2$ are mean centered to standard deviation of 1. Significant results below significance level $\alpha$ = 0.05 are bolded.

| | Estimate | Standard error | Z value | Df | Chisq | Pr (>Chisq) |
|---|---|---|---|---|---|---|
| **Injury treatment** | **-0.10** | **0.10** | **-1.04** | **2** | **99.28** | **<0.001** |
| **Infection treatment** | **-1.00** | **0.10** | **-10.13** | | | |
| Protein | -0.02 | 0.04 | -0.59 | 1 | 0.35 | 0.56 |
| Protein$^2$ | 0.02 | 0.05 | 0.47 | 1 | 0.22 | 0.64 |

**Table S8:** Model summary of a negative binomial model of the effects of protein in larval diet and stress treatments on lifespan of adult flies. Protein and protein$^2$ are mean centered to standard deviation of 1.

| | Estimate | Standard error | Z value | Df | Chisq | Pr (>Chisq) |
|---|---|---|---|---|---|---|
| Intercept | 4.38 | 0.12 | 37.63 | | | |
| Injury treatment | -0.06 | 0.16 | -0.40 | | | |
| Infection treatment | -0.97 | 0.16 | -6.00 | | | |
| Protein | -0.10 | 0.08 | -1.22 | | | |
| Protein$^2$ | 0.05 | 0.10 | 0.49 | | | |
| Injury:Protein | 0.12 | 0.11 | 1.11 | 2 | 1.22 | 0.54 |
| Infection:Protein | 0.06 | 0.11 | 0.56 | | | |
| Injury:Protein$^2$ | -0.04 | 0.13 | -0.28 | 2 | 0.08 | 0.96 |
| Infection:Protein$^2$ | -0.03 | 0.13 | -0.21 | | | |

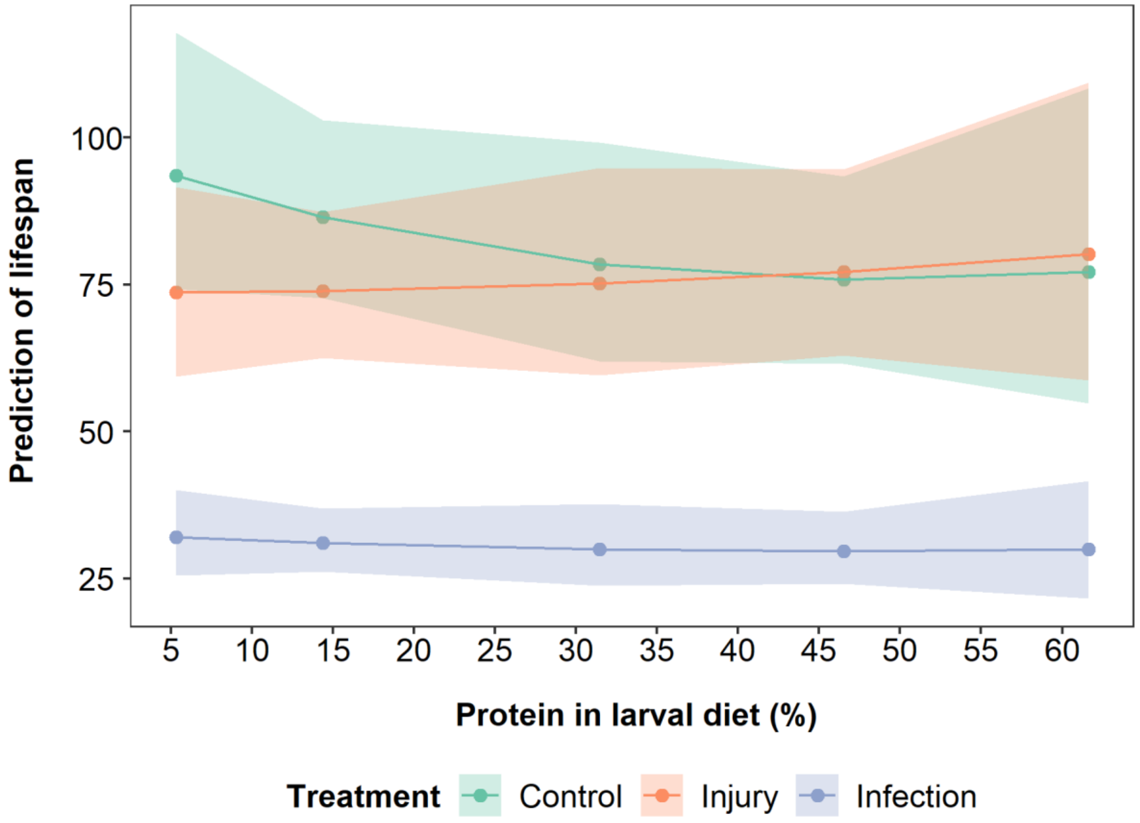


**Figure S7:** Model predictions of the effects of larval protein restriction on adult lifespan of flies infected with a bacterial pathogen (blue data points and lines), injured by a pinprick (orange data points and lines) or with no treatment (green data points and lines). Shaded areas are 95% confidence intervals.

**Table S9:** Model summary of a Cox Proportional Hazard regression model of the effects of protein in larval diet and stress treatments on survival (n = 407, number of deaths = 365, concordance = 0.672, $R^2 = 0.18$). Protein and protein$^2$ are mean centered to standard deviation of 1. Significant results below significance level $\alpha = 0.05$ are bolded.

| | coef | exp(coef) | se(coef) | z | Pr (>\|z\|) |
|---|---|---|---|---|---|
| Injury treatment | 1.15 | 1.15 | 021 | 0.67 | 0.50 |
| **Infection treatment** | **0.92** | **2.50** | **0.21** | **4.32** | **<0.001** |
| Protein | 0.20 | 1.22 | 0.11 | 1.82 | 0.07 |
| Protein$^2$ | -0.19 | 0.83 | 0.12 | -1.54 | 0.12 |
| Injury:Protein | -0.16 | 0.86 | 0.15 | -1.03 | 0.30 |
| Infection:Protein | -0.17 | 0.84 | 0.15 | -1.11 | 0.27 |
| Injury:Protein$^2$ | 0.11 | 1.11 | 0.17 | 0.63 | 0.53 |
| Infection:Protein$^2$ | 0.25 | 1.29 | 0.17 | 1.48 | 0.14 |

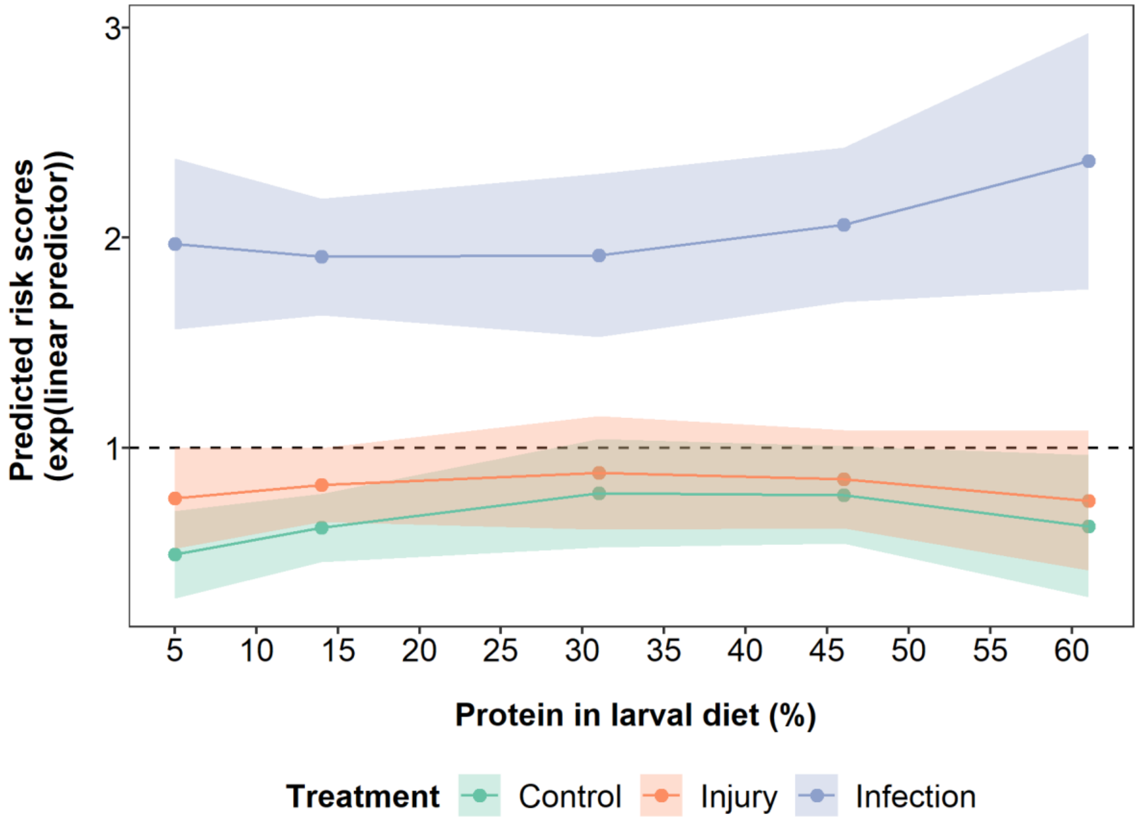


**Figure S8:** Model predictions of the effects of larval protein restriction on survival of flies infected with a bacterial pathogen (blue data points and lines), injured by a pinprick (orange data points and lines) or with no treatment (green data points and lines). $y = 1$ line shows no change in risk ratio, i.e. treatment would have no effect compared to baseline hazard. Shaded areas are 95% confidence intervals.

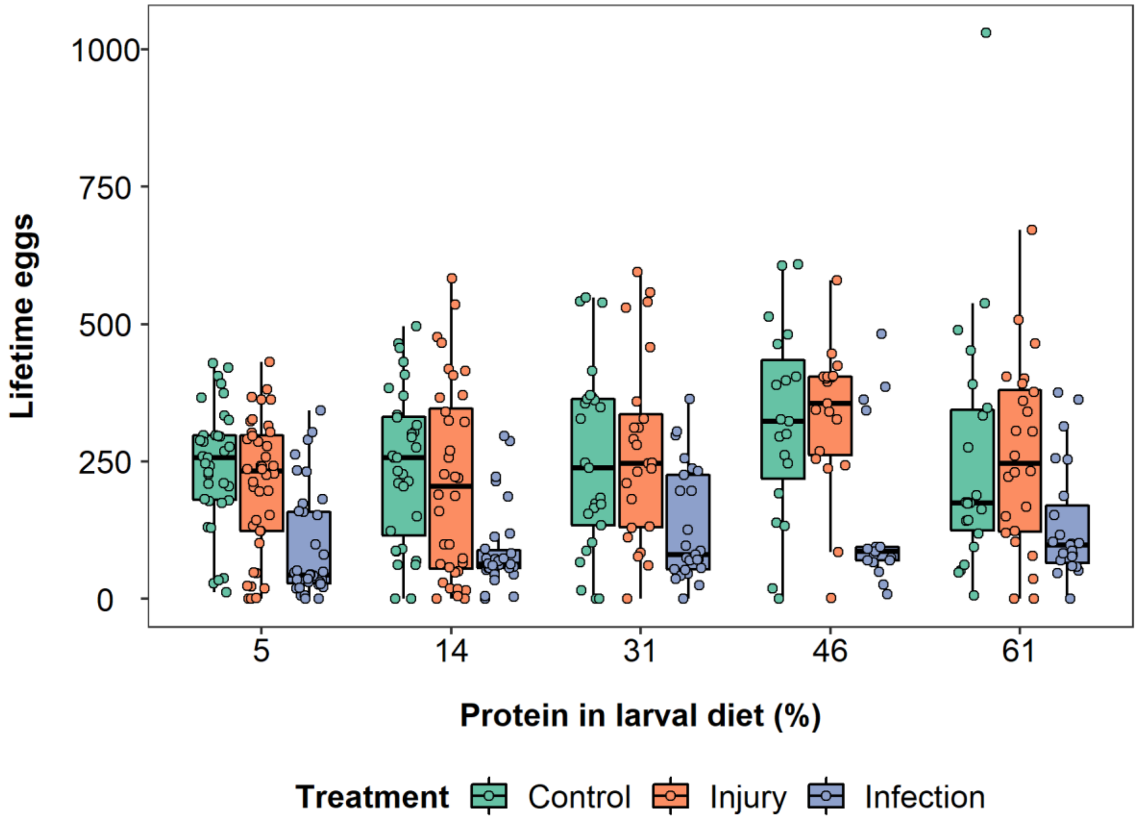


**Figure S9:** Effects of protein in larval diet on the lifetime eggs produced per female (up to day 98) of flies infected with a bacterial pathogen (blue data points and lines), injured by a pinprick (orange data points and lines) or with no treatment (green data points and lines). The lines in the box plots indicates median values (50% quantile), boxes are the interquartile range (25% to 75% quantiles) and whiskers are minimum or maximum quartiles (25% - 1.5 x interquartile range, 75% + 1.5 x interquartile range).

**Table S10:** Summary of main effects parameter estimates and associated LRT test values for a zero-inflated negative binomial model of the effects of protein in larval diet and stress treatments on the total number of eggs produced per fly with lifespan added as a term in the model. The values are from models not including interactions with the specific main effect. Chi-squared and associated p-values are from LRT tests comparing a model with no interactions associated with the main effect to a model with no main effect. Protein protein$^2$ and lifespan are mean centered to standard deviation of 1. Significant results below significance level $\alpha = 0.05$ are bolded.

| | Estimate | Standard error | Z value | Df | Chisq | Pr (>Chisq) |
|---|---|---|---|---|---|---|
| **Injury treatment** | **0.04** | **0.07** | **0.63** | **2** | **8.81** | **0.01** |
| **Infection treatment** | **-0.18** | **0.08** | **-2.30** | | | |
| **Protein** | **0.11** | **0.05** | **2.45** | **1** | **5.73** | **0.02** |
| **Protein$^2$** | **-0.11** | **0.04** | **-3.04** | **1** | **8.01** | **0.005** |

**Table S11:** Model summary of a zero-inflated negative binomial model of the effects of protein in larval diet and stress treatments on the total number of eggs produced per fly. Block and individual ID are added as random effects. Lifespan is added in the model to account for selective disappearance. Protein protein$^2$ and lifespan are mean centered to standard deviation of 1. Significant results below significance level $\alpha = 0.05$ are bolded.

| | Estimate | Standard error | Z value | Df | Chisq | Pr (>Chisq) |
|---|---|---|---|---|---|---|
| Intercept | 5.27 | 0.08 | 64.23 | | | |
| Injury treatment | 0.06 | 0.11 | 0.54 | | | |
| Infection treatment | -0.17 | 0.12 | -1.41 | | | |
| Protein | 0.18 | 0.06 | 3.10 | | | |
| Protein$^2$ | -0.10 | 0.07 | -1.60 | | | |
| **Lifespan** | **0.69** | **0.04** | **19.78** | **1** | **275.77** | **<0.001** |
| Injury:Protein | -0.02 | 0.08 | -0.29 | 2 | 2.39 | 0.30 |
| Infection:Protein | 0.10 | 0.08 | 1.18 | | | |
| Injury:Protein$^2$ | -0.02 | 0.09 | -0.24 | 2 | 0.06 | 0.97 |

**Table S12:** Summary of main effects parameter estimates and associated LRT test values for a zero-inflated negative binomial model of the effects of protein in larval diet and stress treatments on the total number of eggs produced per fly. The values are from models not including interactions with the specific main effect. Chi-squared and associated p-values are from LRT tests comparing a model with no interactions associated with the main effect to a model with no main effect. Protein protein$^2$ and lifespan are mean centered to standard deviation of 1. Significant results below significance level $\alpha = 0.05$ are bolded.

| | Estimate | Standard error | Z value | Df | Chisq | Pr (>Chisq) |
|---|---|---|---|---|---|---|
| **Injury treatment** | **-0.04** | **0.09** | **-0.48** | **2** | **80.68** | **<0.001** |
| **Infection treatment** | **-0.82** | **0.09** | **-8.82** | | | |
| **Protein** | **0.12** | **0.04** | **3.15** | **1** | **5.91** | **0.02** |
| Protein$^2$ | -0.07 | 0.05 | -1.45 | 1 | 2.10 | 0.15 |

**Table S13:** Model summary of a zero-inflated negative binomial model of the effects of protein in larval diet and stress treatments on the total number of eggs produced per fly. Block and individual ID are added as random effects. Protein protein$^2$ and lifespan are mean centered to standard deviation of 1. Significant results below significance level $\alpha = 0.05$ are bolded.

| | Estimate | Standard error | Z value | Df | Chisq | Pr (>Chisq) |
|---|---|---|---|---|---|---|
| Intercept | 5.67 | 0.11 | 50.99 | | | |
| Injury treatment | -0.05 | 0.16 | -0.32 | | | |
| Infection treatment | -0.86 | 0.16 | -5.54 | | | |
| Protein | 0.09 | 0.08 | 1.21 | | | |
| Protein$^2$ | -0.09 | 0.09 | -0.99 | | | |
| Injury:Protein | 0.08 | 0.11 | 0.75 | 2 | 0.98 | 0.61 |
| Infection:Protein | 0.10 | 0.11 | 0.93 | | | |
| Injury:Protein$^2$ | 0.01 | 0.13 | 0.07 | 2 | 0.10 | 0.95 |
| Infection:Protein$^2$ | 0.04 | 0.13 | 0.29 | | | |
| Infection:Protein$^2$ | -0.01 | 0.09 | -0.08 | | | |

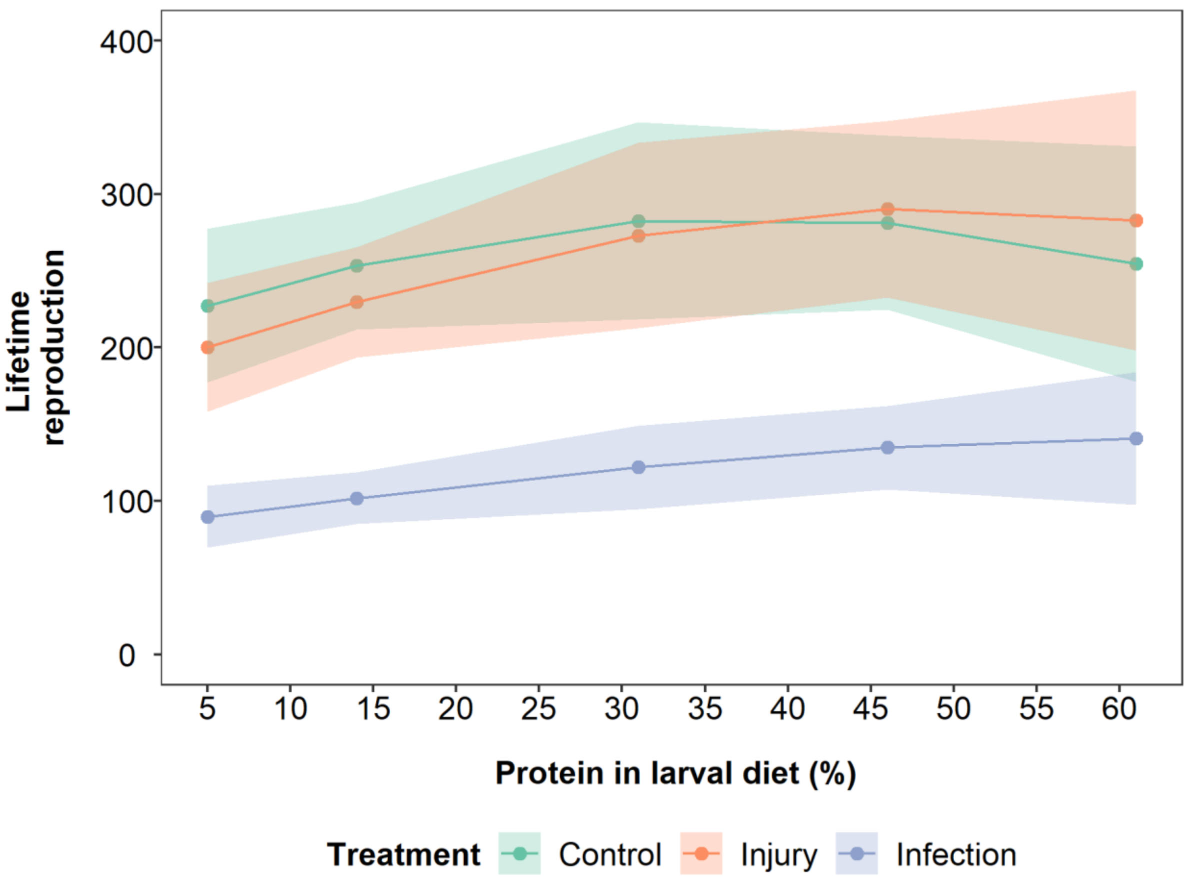


**Figure S10:** Model predictions of the effects of larval protein restriction on lifetime egg production (up to day 98) of flies infected with a bacterial pathogen (blue data points and lines), injured by a pinprick (orange data points and lines) or with no treatment (green data points and lines). Shaded areas are 95% confidence intervals.

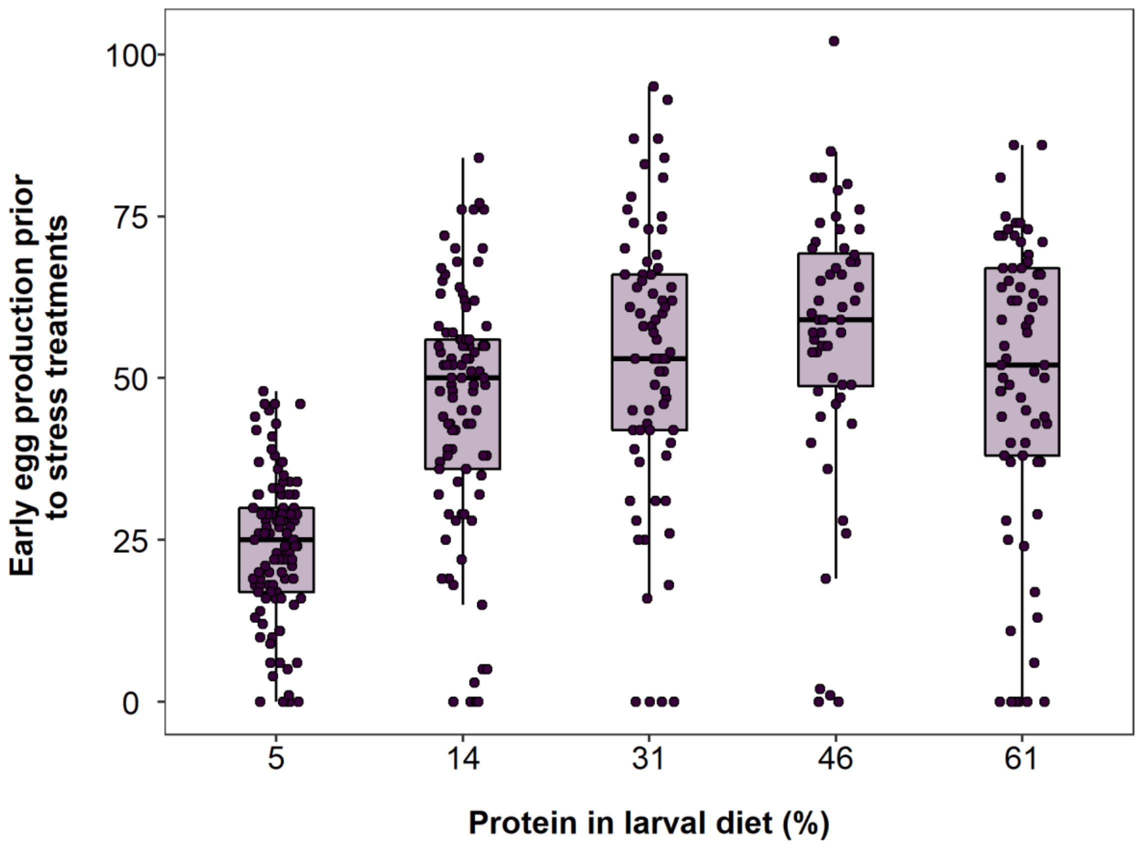


**Figure S11:** Effects of protein in larval diet on the number of eggs produced in the first week before stress treatments. The lines in the box plots indicate median values (50% quantile), boxes are the interquartile range (25% to 75% quantiles) and whiskers are minimum or maximum quartiles (25% - 1.5 x interquartile range, 75% + 1.5 x interquartile range).

**Table S14:** Model summary of a zero-inflated negative binomial model of the effects of protein in larval diet and stress treatments on the total number of eggs produced per fly in the first week. Block and individual ID are added as random effects. Protein protein$^2$ and lifespan are mean centered to standard deviation of 1. Significant results below significance level α = 0.05 are bolded.

| | Estimate | Standard error | Z value | Df | Chisq | Pr (>Chisq) |
|---|---|---|---|---|---|---|
| Intercept | 3.97 | 0.08 | 48.60 | | | |
| Protein | 0.29 | 0.06 | 5.22 | 1 | 3.71 | 0.054 |
| **Protein$^2$** | **-0.20** | **0.04** | **-4.77** | **1** | **22.19** | **<0.001** |

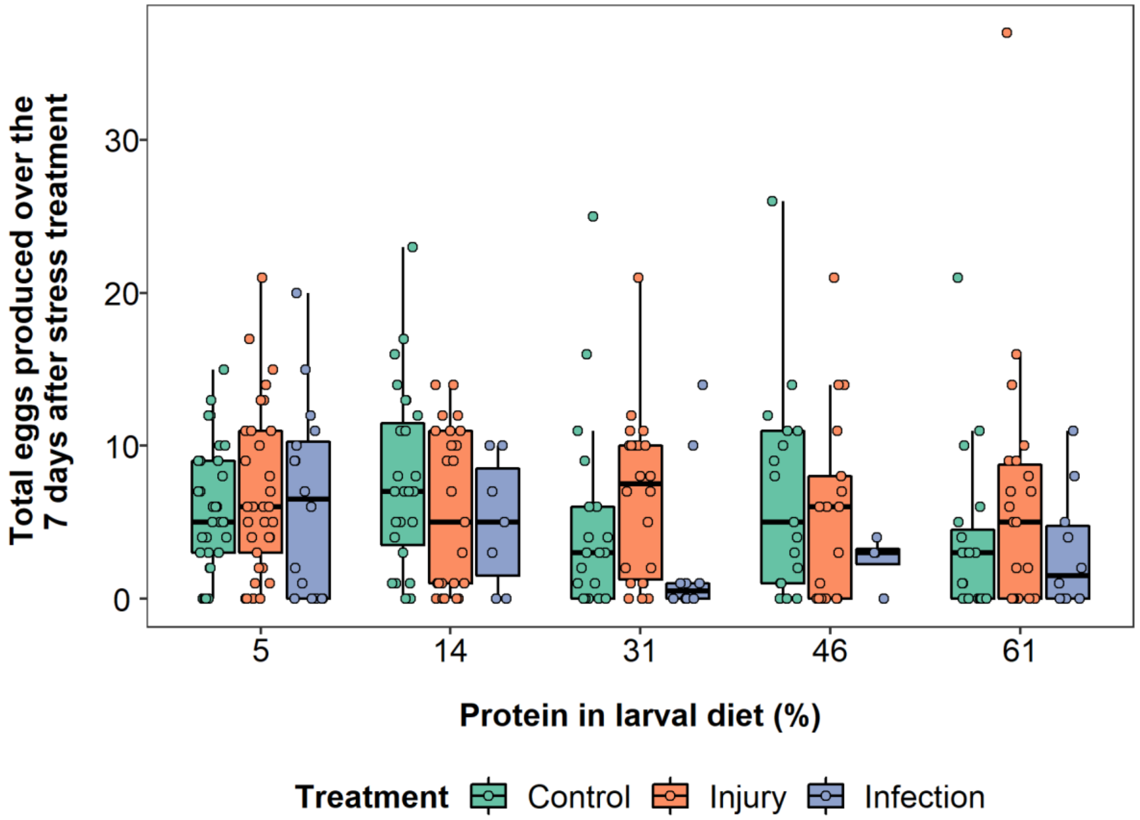


**Figure S12:** Effects of protein in larval diet on total eggs produced over seven days after stress treatment by flies infected with a bacterial pathogen (blue data points and bars), injured by a pinprick (orange data points and bars) or with no treatment (green data points and bars). The lines in the box plots indicate median values (50% quantile), boxes are the interquartile range (25% to 75% quantiles) and whiskers are minimum or maximum quartiles (25% - 1.5 x interquartile range, 75% + 1.5 x interquartile range).

**Table S15:** Summary of main effects parameter estimates and associated LRT test values for a negative binomial model of the effects of protein in larval diet and stress treatments on the total number of eggs produced per fly seven days after stress treatments. The values are from models not including interactions with the specific main effect. Chi-squared and associated p-values are from LRT tests comparing a model with no interactions associated with the main effect to a model with no main effect. Protein and protein$^2$ are mean centered to standard deviation of 1.

| | Estimate | Standard error | Z value | Df | Chisq | Pr (>Chisq) |
|---|---|---|---|---|---|---|
| Injury treatment | 0.12 | 0.12 | 0.99 | 2 | 2.36 | 0.31 |
| Infection treatment | -0.13 | 0.18 | -0.74 | | | |
| Protein | -0.05 | 0.06 | -0.76 | 1 | 0.59 | 0.44 |
| Protein$^2$ | 0.003 | 0.08 | 0.04 | 1 | 0.002 | 0.97 |

**Table S16:** Model summary of a negative binomial model of the effects of protein in larval diet and stress treatments on the total number of eggs produced per fly seven days after stress treatments. Block and individual ID are added as random effects. Protein protein$^2$ and lifespan are mean centered to standard deviation of 1. Block variance = 0.001, standard deviation = 0.02.

| | Estimate | Standard error | Z value | Df | Chisq | Pr (>\|t\|) |
|---|---|---|---|---|---|---|
| Intercept | 2.08 | 0.14 | 14.37 | | | |
| Injury treatment | -0.07 | 0.20 | -0.35 | | | |
| Infection treatment | -0.62 | 0.34 | -1.79 | | | |
| Protein | -0.02 | 0.10 | -0.15 | | | |
| Protein$^2$ | 0.14 | 0.13 | -1.09 | | | |
| Injury:Protein | 0.06 | 0.14 | 0.43 | 2 | 5.57 | 0.06 |
| Infection:Protein | -0.40 | 0.20 | -1.97 | | | |
| Injury:Protein$^2$ | 0.20 | 0.17 | 1.19 | 2 | 2.73 | 0.26 |
| Infection:Protein$^2$ | 0.39 | 0.25 | 1.52 | | | |

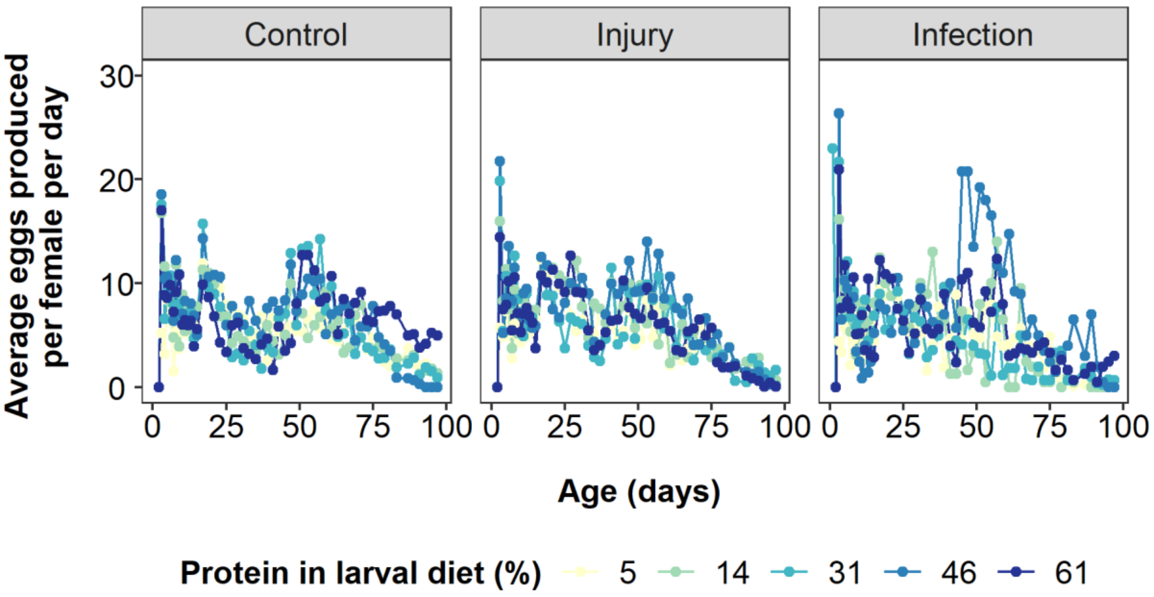


**Figure S13:** Average eggs per day for each stress and larval diet treatments on flies infected with a bacterial pathogen ("Infection"), injured by a pinprick ("Injury") or with no treatment ("Control"). For clarity, associated errors have been removed from the plot.

**Table S17:** Summary of main effects parameter estimates and associated LRT test values for a zero-inflated negative binomial model of the effects of protein in larval diet and stress treatments on the daily number of eggs produced per fly. The values are from models not including interactions with the specific main effect. Chi-squared and associated p-values are from LRT tests comparing a model with no interactions associated with the main effect to a model with no main effect. Protein and protein$^2$ are mean centered to standard deviation of 1. Significant results below significance level $\alpha = 0.05$ are bolded.

| | Estimate | Standard error | Z value | Df | Chisq | Pr (>Chisq) |
|---|---|---|---|---|---|---|
| Injury treatment | -0.02 | 0.04 | -0.56 | 2 | 0.36 | 0.84 |
| Infection treatment | -0.02 | 0.05 | -0.44 | | | |
| Protein | 0.02 | 0.03 | 0.78 | 1 | 0.61 | 0.44 |
| **Protein$^2$** | **-0.12** | **0.03** | **-4.05** | **1** | **12.26** | **0.0005** |
| **Age** | **-0.59** | **0.01** | **-47.22** | **1** | **2175.5** | **<0.001** |
| **Age2** | **-0.07** | **0.01** | **-5.13** | **1** | **26.14** | **<0.001** |

**Table S18:** Summary of two-way interaction estimates and associated LRT test values for a zero-inflated negative binomial model of the effects of protein in larval diet and stress treatments on the daily number of eggs produced per fly. The values are from models not including interactions with the specific main effect. Chi-squared and associated p-values are from LRT tests comparing a model with no interactions associated with the main effect to a model with no main effect. Protein and protein$^2$ are mean centered to standard deviation of 1. Significant results below significance level $\alpha = 0.05$ are bolded.

| | Estimate | Standard error | Z value | Df | Chisq | Pr (>Chisq) |
|---|---|---|---|---|---|---|
| **Protein:Age** | **-0.07** | **0.01** | **-5.52** | **1** | **30.48** | **<0.001** |
| **Protein:Age2** | **0.07** | **0.01** | **4.88** | **1** | **23.94** | **<0.001** |
| **Protein$^2$:Age** | **0.07** | **0.02** | **4.76** | **1** | **22.62** | **<0.001** |

**Table S19:** Model summary of a zero-inflated negative binomial model of the effects of protein in larval diet and stress treatments on the daily number of eggs produced per fly. Lifetime egg counts go up to day 98. Block, individual ID and a value for each row are added as random effects. Protein, protein$^2$, age, age$^2$ and lifespan are mean centered to standard deviation of 1. Significant results below significance level $\alpha = 0.05$ are bolded.

| | Estimate | Standard error | Z value | Df | Chisq | Pr (>Chisq) |
|---|---|---|---|---|---|---|
| Intercept | 1.71 | 0.07 | 26.14 | | | |
| Injury treatment | 0.08 | 0.08 | 1.06 | | | |
| Infection treatment | -0.09 | 0.10 | -0.86 | | | |
| Protein | 0.02 | 0.05 | 0.41 | | | |
| Protein$^2$ | -0.05 | 0.05 | -1.13 | | | |
| Age | -0.60 | 0.03 | -21.74 | | | |
| Age$^2$ | -0.03 | 0.02 | -1.63 | | | |
| Lifespan | 0.02 | 0.02 | 1.41 | 1 | 2.00 | 0.16 |
| Injury:Protein | 0.07 | 0.06 | 1.17 | 2 | 1.61 | 0.45 |
| Infection:Protein | 0.002 | 0.07 | 0.03 | | | |
| Injury:Protein$^2$ | -0.09 | 0.06 | -1.54 | 2 | 2.66 | 0.26 |
| Infection:Protein$^2$ | -0.01 | 0.07 | -0.15 | | | |
| **Injury:Age** | **-0.05** | **0.04** | **-1.18** | **2** | **18.05** | **<0.001** |
| **Infection:Age** | **-0.25** | **0.06** | **-4.27** | | | |
| Injury:Age$^2$ | 0.07 | 0.03 | -2.30 | 2 | 5.90 | 0.052 |
| Infection:Age$^2$ | -0.06 | 0.04 | -1.50 | | | |
| Protein:Age | -0.10 | 0.02 | -4.86 | | | |
| Protein:Age$^2$ | 0.10 | 0.02 | 4.24 | | | |
| Protein$^2$:Age | 0.12 | 0.02 | 5.10 | | | |
| Injury:Protein:Age | 0.05 | 0.03 | 1.64 | 2 | 2.75 | 0.25 |
| Infection:Protein:Age | 0.03 | 0.04 | 0.82 | | | |
| **Injury:Protein:Age$^2$** | **-0.08** | **0.03** | **-2.54** | **2** | **14.41** | **0.001** |
| **Infection:Protein:Age$^2$** | **0.08** | **0.04** | **1.75** | | | |
| **Injury:Protein$^2$:Age** | **-0.10** | **0.03** | **-3.18** | **2** | **11.21** | **0.004** |
| **Infection:Protein$^2$:Age** | **-0.003** | **0.05** | **-0.07** | | | |
